# Integrating Metabolic Networks into Hybrid Bioprocess Models

**DOI:** 10.64898/2026.04.22.720062

**Authors:** Mathias Gotsmy, Gonzalo Guillén-Gosálbez

**Affiliations:** ETH Zürich, Vladimir-Prelog-Weg 1-5, Zürich, 8093, Switzerland

**Keywords:** bioprocess optimization, hybrid modeling, surrogate flux balance analysis, symbolic regression, neural differential equations

## Abstract

The optimization and control of bioprocesses require robust *in silico* models that can accurately capture the complex and dynamic behavior of living cells. While hybrid models that combine machine learning with mechanistic equations have emerged as a powerful tools, they often require relatively large datasets and might yield inconsistent predictions that violate the stoichiometry of metabolism.

In this study, we introduce `FBA-Hyb`, a multi-scale hybrid modeling framework that tightly integrates genome-scale metabolic networks via flux balance analysis (FBA) into its architecture. In our `FBA-Hyb` framework, artificial neural networks predict key FBA inputs (substrate uptake rates and FBA objectives) while a surrogate FBA module translates them into the metabolic fluxes that govern the bioprocess. A key novelty is that the FBA optimization step is replaced by a surrogate generated with symbolic regression, which encapsulates the FBA model into a compact analytical expression. This allows easy backpropagation through the integration of the neural controlled differential equationbased `FBA-Hyb` bioprocess model.

We validated `FBA-Hyb` against a standard hybrid model (`Std-Hyb`) using two *Escherichia coli* fedbatch case studies. In the first study, `FBA-Hyb` achieved a 42% average improvement in predictive accuracy (*R*^2^) during a leave-one-process-out cross validation. Crucially, `FBA-Hyb` maintains strict stoichiometric feasibility even during extrapolation. Meanwhile, an alternative approach based on standard architectures leads to stoichiometrically inconsistent solutions in 22% of the cases analyzed. In the second case study, we demonstrate how `FBA-Hyb` effectively simulates unmeasured chemical species and discovers a metabolic shift in sulfate-limited regimes during bioprocessing.

By providing a modular, biologically consistent, and computationally efficient architecture, `FBA-Hyb` offers a robust foundation for the next generation of bioprocess models and sustainable process optimization.

**Highlights:**

- `FBA-Hyb` integrates flux balance analysis (FBA) into hybrid bioprocess models.
- Symbolic regression discovers a simple closed-form FBA surrogate model.
- The FBA surrogate ensures accurate reaction stoichiometry.
- A neural network predicting the FBA objective keeps the model flexible.
- `FBA-Hyb` has superior capabilities and accuracy compared to the current standard.

**Graphical Abstract:** 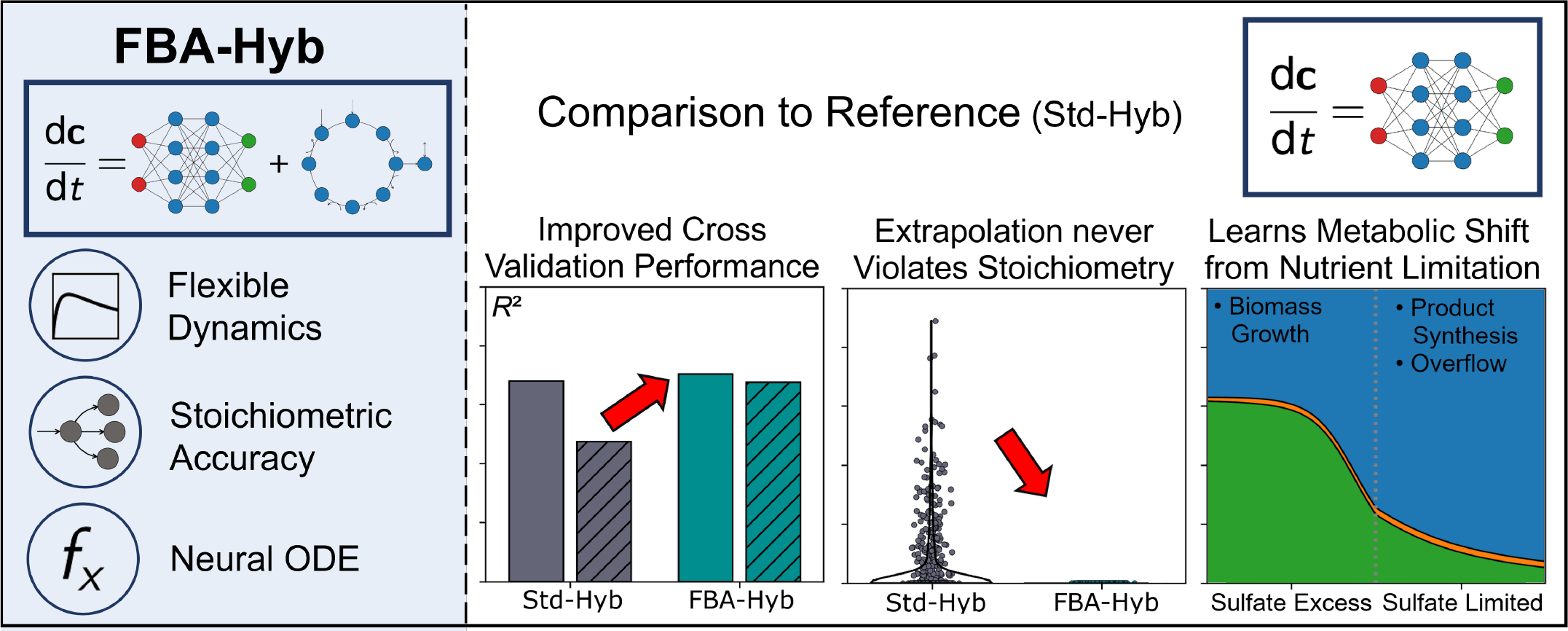

## 1. Introduction

Bioprocess have become an essential tool in biotechnology, enabling the large-scale production of organic compounds from commodity chemicals to life-saving pharmaceuticals [1]. Moreover, bioprocesses offer pathways to de-fossilize the chemical industry by providing renewable routes to compounds traditionally derived from petroleum [2]. Unlike conventional chemical processes, bioprocesses rely on living cells with many of the essential product synthesis steps occurring intracellularly [3]. This makes them uniquely powerful but also challenging: cellular systems are complex, dynamic, and environment-dependent, making their behavior difficult to predict and control.

Addressing these challenges requires robust and accurate *in silico* bioprocess models. Bioprocess models predict cellular responses and bioprocess outcomes such as product quantity and quality from process control parameters. The obtained insights can guide scientists during process development and monitoring. Bioprocess models allow bioprocesses to be optimized faster, products to meet quality standards more reliably, and development cycles to become more sustainable and cost-efficient [4]. Consequently, the development of robust and accurate bioprocess models is a fundamental basis for the next generation of digital and sustainable biotechnology.

Over the past decades, a constant stream of new bioprocess modeling approaches has been published, reflecting ongoing advances in both biological understanding and computational techniques [5]. Many previous studies on bioprocess modeling focused on building kinetic representations of the system based on well-established formalisms [6]. Increasingly, these models are being enriched through integration with more granular approaches such as flux balance analysis (FBA). The challenge here is to integrate both modeling frameworks, originally developed by different communities and which often serve different purposes, within a holistic approach to more accurately describe the bioprocess’ behavior. We shall elaborate further on this later in the article.

### 1.1. Kinetic modeling of bioprocesses

Kinetic bioprocess models are developed by generalizing the (semi-)empirical equations of enzyme kinetics to whole cells [7]. Despite the simplification of lumping many chemical reactions into relatively few equations, kinetics-based bioprocess models have proven highly effective in practice [8]. In addition to being comparatively robust to overfitting, one major advantage is their ability to capture changes in cell kinetics (e.g. over time or due to shifts in pH or temperature) [9, 10]. However, despite performing well on specific processes, the development of such models is time-intensive as the impact of the relevant control variables on the kinetics needs to be determined which may take many experiments and iterations [6].

### 1.2. Dynamic flux balance analysis

FBA requires three inputs: (i) a genome-scale metabolic model (GSMM) which includes all relevant metabolic reactions of a cell, (ii) bounds on the respective fluxes (i.e. metabolic reaction rates), and (iii) a user-defined objective function. Using these inputs, FBA calculates the fluxes that optimize the objective reaction by solving a linear program (LP) [11]. Dynamic FBA (dFBA) extends the steady-state solutions of FBA to (bioprocess) time-series by integrating predicted fluxes over time [12]. Dynamics are typically introduced by modeling substrate uptake fluxes with kinetic equations [13]. However, FBA itself is an LP, so its integration into models at higher scales naturally results in a bilevel optimization problem, where the inner problem is the FBA while the outer problem, for example during model building, seeks to minimize the mismatch of the model predictions with experimental data. Such bilevel problems are known to be very hard to solve, particularly when nonconvexities are present [14, 15].

### 1.3. Hybrid bioprocess models

Hybrid models embed machine learning (ML) components into mechanistic frameworks, allowing unknown system dynamics to be learned from data while preserving established process knowledge [16]. This synergy has led to models that are both accurate and robust, even in data-limited settings, and has stimulated methodological innovation in bioprocess modeling [16]. Various ML architectures have been applied in this context, including (deep) artificial neural networks (ANNs) [17, 18], Gaussian processes (GPs) [19, 20], physics-informed neural networks (PINNs) [21], and long short-term memory (LSTM) networks [22].

Hybrid bioprocess models typically perform well when modeling relationships involving less established control variables for which mechanistic insights are limited [16]. However, compared to bioprocess models based on detailed metabolic network or enzyme kinetic structures, hybrid approaches still require relatively large, high-quality datasets to achieve reliable performance, which are not always available.

### 1.4. Including FBA in hybrid models

Originally, FBA-based and hybrid bioprocess models were largely developed independently by different research communities, despite addressing related questions in bioprocess modeling. As a result, their combination has historically been limited.

Recently, however, there has been growing interest in integrating ML with FBA, and initial studies have begun to explore this direction. For instance, ML has been used to improve FBA accuracy by predicting growth medium-specific flux bounds for *E. coli* and *P. putida* GSMMs [23]. Similarly, ML-predicted flux bounds were shown to enhance intracellular flux predictions obtained via FBA [24]. In another study, enzyme-constrained FBA has been employed to analyze hybrid model predictions after parameter fitting, for uncertainty estimation in mammalian fed-batch bioprocesses [25]. More recently, Richelle et al. proposed a framework in which FBA and a hybrid model are coupled in parallel to improve bioprocess predictions [26].

While these studies highlight the potential of combining FBA and ML, a general and systematic integration framework is still lacking. Several limitations contribute to this gap. First, current approaches that use ML to enhance FBA predictions typically operate in static settings, where flux bounds are predicted for a single condition [23, 24]. However, metabolic fluxes and uptake rates are inherently dynamic during bioprocesses, suggesting that incorporating time-dependent behavior could further improve predictive performance [27]. Second, when FBA is used only for *post hoc* analysis or coupled externally, hybrid models cannot directly learn from FBA-derived constraints. A key technical challenge is that FBA is formulated as a LP, which complicates its integration into end-to-end ML frameworks that rely on gradient-based optimization and backpropagation [28].

One promising direction to address these challenges is the use of surrogate models for FBA. However, this remains an underexplored area. So far, surrogate FBA models have primarily relied on ANNs [29, 23], while alternative approaches such as symbolic regression (SR) have received little attention. SR aims to discover explicit symbolic expressions from data by searching the space of mathematical equations to identify models that best capture the underlying relationships [30]. Due to its interpretability and flexibility, it has recently gained traction as a surrogate modeling approach in (bio)chemical engineering [31, 32]. However, SR has not yet been applied for creating FBA surrogate models.

### 1.5. This study

In this study, we introduce `FBA-Hyb`, a novel framework for combining FBA predictions with hybrid modeling that explicitly addresses the limitations of previous approaches. The key novelty of `FBA-Hyb` lies in its multi-scale architecture, which is designed to preserve the flexibility of hybrid modeling while enforcing the mechanistic constraints of cellular metabolism.

The proposed `FBA-Hyb` architecture introduces two key design choices. First, the FBA linear program is replaced by a surrogate FBA model that is built *a priori* using SR, yielding a highly accurate and fully differentiable symbolic representation of the FBA solution. Second, instead of predicting all metabolic rates directly, the ANN component of the hybrid model predicts a time-, state-, and control-dependent FBA objective reaction, along with the carbon substrate uptake flux. This provides the necessary inputs to evaluate the surrogate FBA model, which in turn computes a stoichiometrically consistent set of metabolic fluxes that can be integrated into the bioprocess model.

Because the surrogate FBA is expressed as a symbolic equation, gradients can be propagated through it, enabling end-to-end training. This allows the hybrid model to learn how the predicted FBA inputs influence metabolic fluxes and to dynamically recalibrate these inputs during optimization.

We evaluate `FBA-Hyb` using two *E. coli* fed-batch case studies and demonstrate that it substantially improves predictive accuracy during cross-validation compared to a standard hybrid model without a surrogate FBA module. In addition, the framework captures the dynamics of unmeasured variables, correctly learning and predicting sulfate uptake fluxes and identifying a metabolic shift caused by sulfate limitation.

## 2. Methods

The study consisted of three main steps (Fig. 1). First, two hybrid model architectures were defined: a reference hybrid model (`Std-Hyb`) and the proposed `FBA-Hyb`, which integrates a surrogate FBA model into the hybrid model. Second, both architectures were trained and tuned in a leaveone-process-out cross-validation (LOPO CV). Third, the resulting LOPO CV models were combined into ensembles, which were used to predict an unseen test set and for the downstream performance analysis.

**Figure 1:**
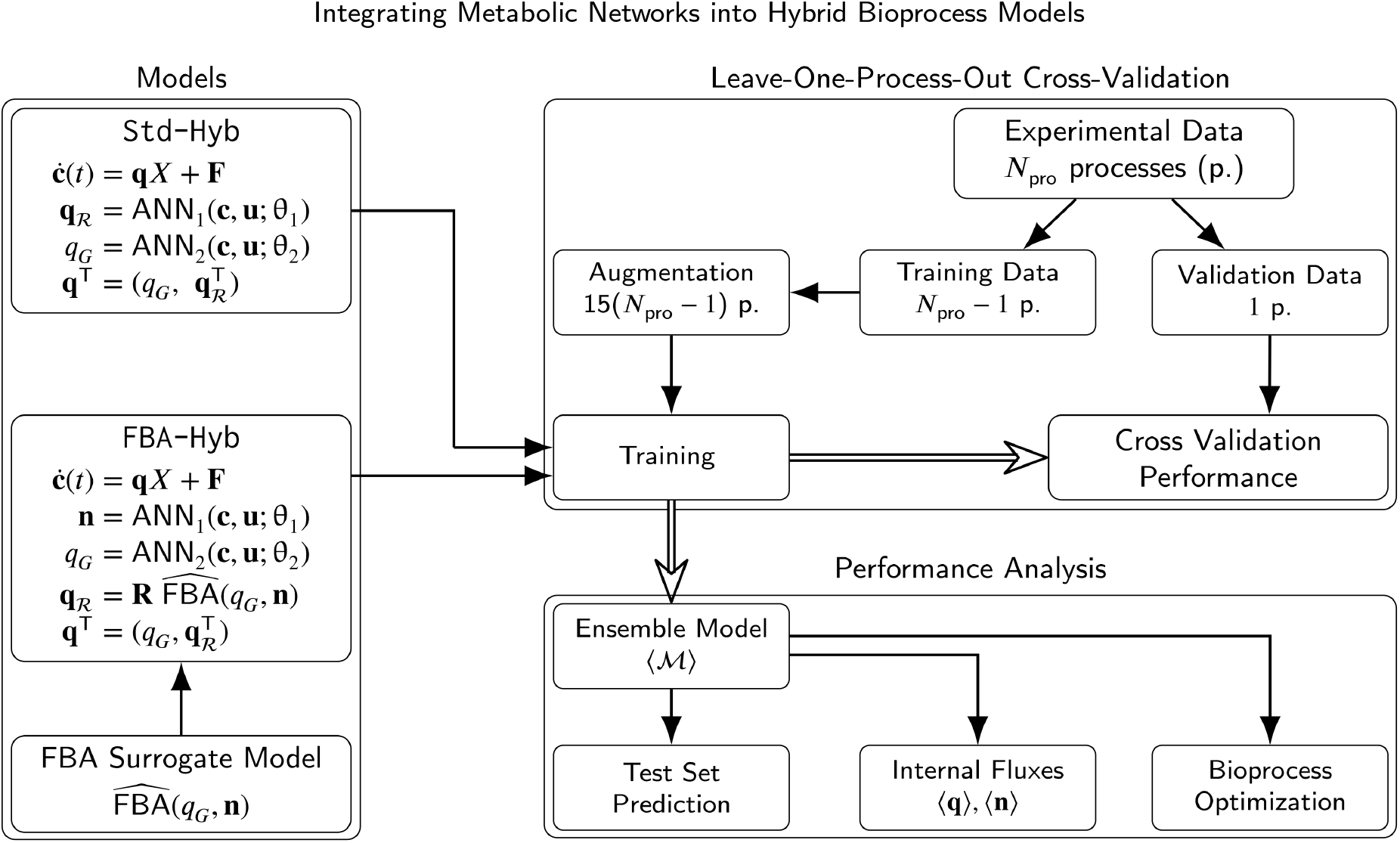
Overview of the modeling and validation pipeline. `Std-Hyb` and `FBA-Hyb` were evaluated in a leave-one-process-out cross-validation (LOPO CV) over all *N*_pro_ process splits. The resulting models were combined into ensembles ⟨ℳ⟩ for the test set prediction, the analysis of internal fluxes, and bioprocess optimization. Double-lined arrows indicate steps that combine the results of all *N*_pro_ LOPO CV models.

### 2.1. The bioprocess controlled ODE

To derive our approach, we consider a general bioprocess controlled ordinary differential equation (ODE) that aims to describe relevant concentrations (i.e., state variables, **c**) in a bioreactor. Typically, they consist of at least substrate (*G*), biomass (*X*), and product (*P*) but may be extended according to the specific bioprocess requirements; such that **c** = (*G, X, P* , …)^T^. The accompanying controlled ODE then reads

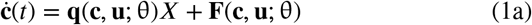

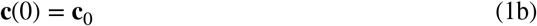

where **q** = (*q*_*G*_, *q*_*X*_, *q*_*P*_ , …)^T^ are the specific metabolic flux rates (i.e., rates of uptake and synthesis of chemicals by the cells), **F** are dilution and feeding terms, and **c**_0_ are initial concentrations. Both, **q** and **F** may depend on dynamic state variables (**c**), dynamic controls (**u**), and constant parameters (θ). Depending on the model architecture, the right-handside of Eq. 1a may be a set of semi-empirical (mechanistic) equations, machine learning modules, or a combination of both: a hybrid model.

In the following sections the mathematical structure of two hybrid models `Std-Hyb` and `FBA-Hyb` is presented. `Std-Hyb` represents a reference hybrid model similar to what has been proposed before, whereas `FBA-Hyb` seamlessly integrates FBA with ANNs into a mechanistic backbone.

### 2.2. Reference hybrid bioprocess model

Here we describe a hybrid bioprocess model in its most common and generalized form [17, 33], from now on referred to as `Std-Hyb` (i.e., the standard approach). This model will be later used for comparison purposes. The two terms of the right hand side of Eq. 1a are modeled as neural controlled ODEs,

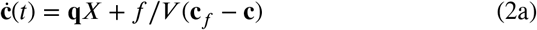

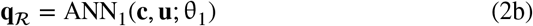

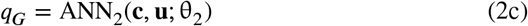

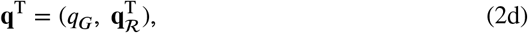

where **q**_*ℛ*_ = (*q*_*X*_, *q*_*P*_ , …)^T^ are all for the bioprocess relevant metabolic fluxes other than the substrate uptake rate (*q*_*G*_). The separation of **q** into *q*_*G*_ and **q**_*ℛ*_ and their prediction by two distinct ANNs (Eqs. 2b and 2c) is introduced to facilitate customization. Note that ANN_2_ can easily be replaced by an analytical equation, if necessary. Many fed-batch processes are substrate-limited and an analytical equation for *q*_*G*_ can be derived [34]. The change of concentrations due to feeding (Eq. 2a) is described by *f*/*V* (**c**_*f*_ − **c**) where *f* is the feed rate and **c**_*f*_ are the feed concentrations. Various widths and depths of the ANNs are reported in literature and need to be adjusted to the case study at hand [17, 33]. We report them in Section 2.9: Case Studies, individually.

### 2.3. FBA hybrid bioprocess model

We propose an alternative hybrid model that integrates FBA via surrogate modeling to enhance its predictive capabilities, as described next. The novel `FBA-Hyb` approach presented here consists of two modeling steps:

1. Creation of an FBA surrogate model that predicts optimal metabolic flux distributions as a function of the carbon substrate uptake rate (*q*_*G*_) and the FBA objective weights (**n**).
2. Creation of a hybrid model of the form of an neural controlled ODE. There the ANNs dynamically predict inputs for the surrogate FBA and the resulting outputs are used to calculate the right hand side of the differential equations.

Recall that the main novelty here lies in embedding the FBA surrogate within the hybrid bioprocess model ODE for easy backpropagation during training. Both modeling steps are explained in more detail the following sections.

#### 2.3.1. FBA surrogate model generation

The well-established parsimonious FBA method allows the estimation of all intra- and extracellular metabolic fluxes (**v**) of a cell from a GSMM [35]. However, parsimonious FBA was developed for steady-state predictions. To account for the typically varying fluxes observed in a bioprocess [34], two strategies have been explored: one accomplish this by either (i) adapting the flux bounds (**v**_min_ and **v**_max_) or (ii) modifying the objective reaction (*v*_obj_) [11]. In this work, we focus on a mixed approach where we vary the flux bounds of the main carbon substrate uptake rate (*v*_*G*,min_ = *v*_*G*_ = *v*_*G*,max_) and the composition of the objective reaction (*v*_obj_).

Different objective reactions were constructed following the structure

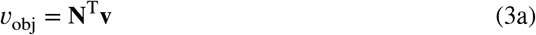

where **N** is a weight vector. In practice the weight vector is 0 for most fluxes, therefore, we refer to the non-zero weight vector as **n** from now on. Rearranging the rows of **v** gives

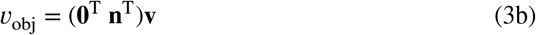

The selection of fluxes included in **n** as well as the identification of the relevant carbon substrate uptake flux (*v*_*G*_) depend on the specific bioprocess under consideration. In practice, **n** includes all relevant metabolic fates a carbon substrate molecule can have in a bioprocess. For example, for both case studies presented here, the vector **n** consisted of weights for three central reactions: the biomass production, the product synthesis, and the ATP maintenance (for details see Sec. 2.9).

To generate training data for FBA surrogate modeling, values for **n** as well as the carbon substrate update rate *v*_*G*_ were then Latin hypercube-sampled 10,000 times for realistic ranges. After this FBA input data sampling, the output data was calculated by optimizing a parsimonious FBA for all 10,000 conditions via solving following bilevel LP optimization problem. Recall that a parsimonious FBA is a bilevel optimization problem that after optimizing the objective reaction also minimizes the sum of all fluxes to ensure uniqueness of the flux distribution [35].

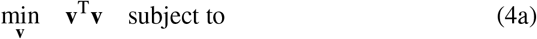

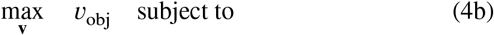

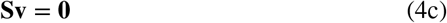

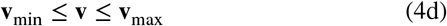

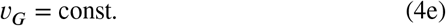

After input (*v*_*G*_, **n**) and output (**v**) data generation, the resulting mapping of the FBA problem is denoted by

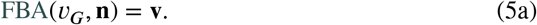

From the full flux vector (**v**), the subvector **q**_*ε*_ ⊂ **v** containing the relevant exchange fluxes (ε) of the FBA model (i.e., uptake and production fluxes of the cell) was extracted and used as target output for surrogate model construction.

Surrogate models are simplified and computationally efficient approximations of more complex models or mappings. For example, the parsimonious FBA method (Eq. 5a), which itself requires solving an optimization problem, can be replaced by a symbolic surrogate that is evaluated as a closed-form function. This significantly reduces computational cost during bioprocess model training and, importantly, enables gradient-based optimization via backpropagation.

To construct such a surrogate for FBA, we employed symbolic regression (SR). Given the generated training data and a set of basic operators (+, -, *, /), the SR algorithm systematically explored the space of mathematical expressions to identify explicit and interpretable equations. These equations approximate the mapping from inputs (*v*_*G*_, **n**) to outputs (**q**_*ε*_) by selecting functions that best reproduce the observed input-output relationships [30]. Initially, one SR run (using `PySR` [30]) was performed for each output of **q**_*ε*_ and, subsequently, the parameters of the found equations were fitted with least-squares minimization [36] resulting in the FBA surrogate model

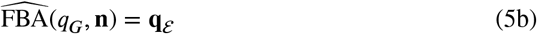

where **q**_*ε*_ includes the surrogate modeling error. We will show that this error is neglible in practice.

#### 2.3.2. Neural controlled differential equation

In our new approach, `FBA-Hyb`, the specific metabolic fluxes (**q**) are not directly calculated from ANNs, but rather from the FBA surrogate model established in previous section. The right hand side of Eq. 1a is modeled as

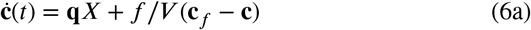

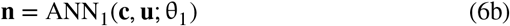

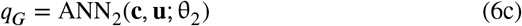

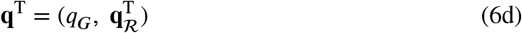

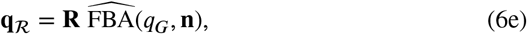

where the reduction matrix **R** ∈ {0 1} ^*ℛ*^ × ^*ε*^ selects the exchange fluxes of the surrogate model FBA that are relevant for the bioprocess at hand, such that **q**_*ℛ*_ = **R q**_*ε*_ is a subvector of **q**_*ε*_. The ANN width and depth were adjusted to the specific case studies, individually, and the ANNs were trained on the respective experimental data of the case studies (Sec. 2.6).

### 2.4. Experimental data processing

Due to sampling of the bioprocess reactor, discrete volume changes arise. To avoid these steps in the state variables, volume normalization is performed as suggested in [6]. A more detailed explanation is given in the supplementary Sec. S1.1. To get continuous values from the presented data, splines were fitted through the measured state and control variables [6].

### 2.5. Data augmentation

We augmented experimental data by sampling values from the fitted splines. For each process, we uniformly sampled 7 timepoints to yield a set of ordered observations (*T*). Spline-smoothed *G, P* , and *X* at these times were independently multiplied by Gaussian noise and clipped at zero. The same procedure was done for initial concentrations (**c**_0_). All control variables were duplicated unchanged. The procedure was repeated 15 times producing 15 augmented runs per experiment. The total set of augmented data (*A*) subsequently was used for model training.

### 2.6. ANN training

Training of the ANNs of `Std-Hyb` and `FBA-Hyb` was done by numerical integration of the respective controlled ODEs and backpropagating the loss through the integration steps. Models were trained on augmented trajectories where the controlled ODE integration was started at the first randomly sampled timepoint. For experimental data prediction (i.e., validation) we integrate from *t* = 0 to *t* = *t*_end_. To ensure that all variables remained in ℝ^+^ and to promote stable training behavior, the input data to the ANNs were mean-scaled (*s*(⋅)), and the corresponding inverse scaling was applied to the outputs to recover physical units required for the mass balance equations (Eqs. 2a and 6a). The scalers were fitted from the measured experimental process data (ANN input) and the generated FBA training data (ANN output). Training minimized a composite objective comprising of the mean absolute error (MAE) of measured and scaled state variables (*G, P* , and *X*) plus a non-negativity penalty,

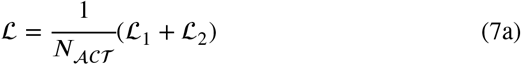

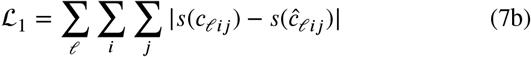

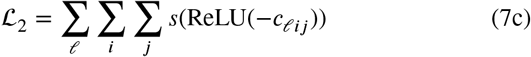

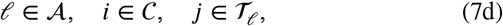

where *A* is the set of augmented processes, *C* is the set of measured state variables, *T*_*l*_ is the set of time points per process, *N*_*ACT*_ is the number of data points in the training data set, and *s* is the scaling function. *c*_*lij*_ are predicted concentrations and ĉ_*lij*_ are augmented experimental values. We used mini-batches of processes, AdamW with weight decay and gradient clipping to stabilize the gradients [37], and early stopping based on validation loss of *P* and *X*.

### 2.7. Cross validation

The `Std-Hyb` and `FBA-Hyb` model configurations and hyperparameters were selected based on a LOPO CV (for details see Sup. Sec. S1.2). For *N*_pro_ measured process time series, *N*_pro_ − 1 processes were used for training and the remaining process for validation. This procedure was repeated *N*_pro_ times, such that each process was used for validation exactly once, resulting in *N*_pro_ trained models per model architecture. The prediction performance was quantified by the normalized mean absolute error (NMAE) and the *R*^2^,

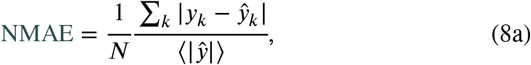

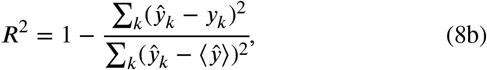

for any predicted *y*_*k*_ and observed quantity ŷ_*k*_, where ⟨⋅⟩ indicates the mean over *N* points and *k* ∈ {1, …, *N*}.

### 2.8. Ensemble predictions

To obtain a single prediction from the *N*_pro_ models of the LOPO CV, we combined them into ensembles,

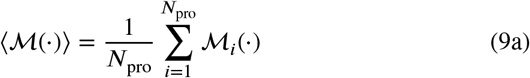

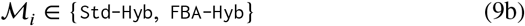

where ℳ_*i*_ is a trained model predictor function of either `Std-Hyb` or `FBA-Hyb`. All analyses following the LOPO CV were performed with these ensembles (Fig. 1). First, the ensembles were validated on a hold-out test set of unseen processes, which was not used for model building, using the same NMAE and *R*^2^ metrics as above. Second, the ensembles were used to analyze the predicted internal fluxes (⟨**q**⟩) and objective weights (⟨**n**⟩), including their stoichiometric feasibility. Third, the ensembles were used to predict process outcomes for conditions that were not part of the training data (i.e., process optimization scenarios).

### 2.9. Case studies

The described hybrid models, `Std-Hyb` and `FBA-Hyb`, were tested on two separate *E. coli* fed-batch process case studies from literature: the production of protein L [6] and the production of plasmid DNA (pDNA) [34] which from now on we will abbreviate as PROL and SLIM, respectively.

In both case studies the non-zero FBA objective weight vector (**n**) was defined as

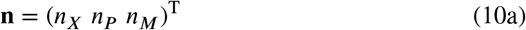

where *n*_*X*_, *n*_*P*_ , and *n*_*M*_ are the objective function weights for the biomass production (*q*_*X*_), product synthesis (*q*_*P*_), and ATP maintenance (i.e., the hydrolysis of ATP to ADP, *q*_*M*_) fluxes of the case studies’ GSMMs, respectively. Relevant exchange fluxes for surrogate FBA model fitting comprised

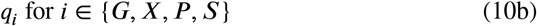

corresponding to the synthesis of biomass, product, and the uptake of SO^2−^, respectively. In both case studies, the product accumula^4^ted internally, therefore, the experimentally measured biomass 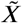 was modeled as

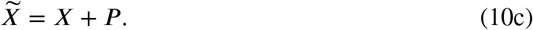

Modeling choices that differed between the two case studies are discussed in the following sections.

### 2.9.1. Protein L production

In the original experimental study, the authors used a transformed *E. coli* BL21(DE3) strain to produce protein L in a fed-batch bioprocess with glycerol as carbon substrate [38]. A full factorial design with two control variables, temperature (*T*) and exponential feed rate (*µ*_*f*_ such that *f*_*G*_(*t*) = *f*_*G*_(0) exp(*µ*_*f*_ *t*)) was conducted [38]. Additionally to the controlled carbon substrate feed (*f*_*G*_) an unknown base feed (*f*_*B*_) needed to be added to the reactor such that

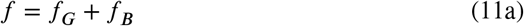

for Eqs. 2a, 6a, and S4a. A linear model was used for its estimation

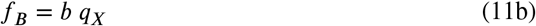

where the parameter *k* was learned during training. A width 16 and depth of 2 was chosen for ANN_1_ and a width 8 and depth 2 for ANN_2_. Softplus were used as activation functions, except for the last layer, where we used

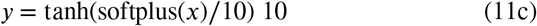

to allow only positive outputs while also constraining the maximum output value for better convergence [28].

The control variables of the process where set as

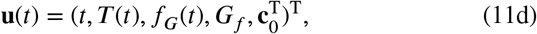

i.e., the time (*t*), the temperature (*T*), the carbon substrate feed (*f*_*G*_), the glycerol concentration in the feed medium (*G*_*f*_), and the initial concentrations (**c**_0_), respectively. *G*_*f*_ and **c**_0_ were constant over one process but changed for different process runs.

### 2.9.2. Plasmid DNA production

In the original pDNA production study, the authors used the *E. coli* strain JM108 in fed-batch bioprocesses with a constant feed rate. Two conditions were tested with multiple replicates: one where sulfate was present in excess (three replicates), and one where sulfate depleted at 21 h into the feed stage (six replicates).

As no base feed was recorded, the equation reduces to

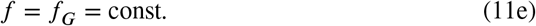

During data augmentation, *t* = 0 was added to all sampled time point sets (*T*) as this is the only time where 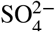 concentrations are known. We chose a width of 64 and depth of 3 for ANN_1_ with softplus activation functions and a custom activation_1_function (Eq. 11c) for the last layer to ensure positive outputs and good convergence.

As in the experimental data the concentration of the carbon substrate (in this case glucose, *G*) was 0 for every timepoint, we replaced the ANN_2_ of `Std-Hyb` and `FBA-Hyb` (Eqs. 2c, 6c) with the analytical equations for the calculation of *q*_*G*_ under steady-state conditions [34],

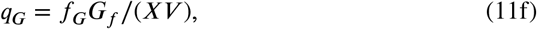

where *G*_*f*_ is the substrate concentration in the feed medium.

The control variables of the process where

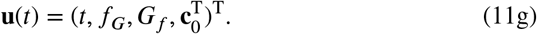

Values of *f*_*G*_, *G*_*f*_ , and **c**_0_ were constant over one process but changed for different process runs.

### 2.10. Implementation

Neural controlled ODEs of `Std-Hyb` and `FBA-Hyb` were trained using `Jax` [39], `Diffrax` [28], and `Equinox` [40]. (Parsimonious) FBA results were calculated using `CobraPy` [41]. Symbolic regression was performed with the `PySR` package [30]. Least squares minimization was done with `SciPy` [36]. All computations were performed on a Windows 11 machine equipped with an AMD Ryzen 7 PRO 8840U CPU (3.30 GHz) and 32 GB of RAM.

### 2.11. Data and code availability

Data and code accompanying this study can be found on GitHub: https://github.com/Gotsmy/FBA-Hyb.

## 3. Results

### 3.1. FBA surrogates are accurate

In order to avoid optimizing an FBA linear program for every step of the numerical integration of the `FBA-Hyb` controlled ODE and to simplify backpropagation, a surrogate FBA model was built (Methods Sec. 2.3.1). By varying the objective weights (**n**) and substrate uptake rate (*q*_*G*_), we generated a parsimonious FBA dataset used to train a compact analytical expression via symbolic regression (SR). Interestingly, the discovered symbolic equations for each output in **q**_*ε*_ converged to the same form:

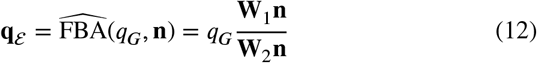

where **W**_1_ and **W**_2_ ∈ ℝ^|*ε*|^×|**n**| are parameter matrices and the division is performed element-wise. Evaluation on both, PROL and SLIM, case studies confirmed the high precision of the surrogate with a validation *R*^2^ of ≈ 1.000 for all components of **q** (Sup. Fig. S1 and S2). The NMAE was ≤ 8.3 ⋅ 10^−5^ and ≤ 5.9 ⋅ 10^−4^ for PROL and SLIM, respectively. Parameters for **W**_1_ and **W**_2_ are reported in Sup. Tab. S1 and S2.

### 3.2. FBA-Hyb outperforms reference models

Fig. 2A compares three distinct modeling approaches (`Std-Hyb`, `FBA-Hyb`, and a literature reference, `OptFed`) on the PROL case study. Their predictive performance was evaluated via leave-one-process-out cross-validation (LOPO CV) and quantified by the coefficient of determination (*R*^2^). Here, *R*^2^ values are reported separately for *P* and *X*, as their prediction difficulty differs (*R*^2^ for biomass is overall lower in this case study). However, since both states are coupled via the shared substrate and metabolic fluxes in the bioprocess ODE, we assess overall model performance by their average. Compared to the `Std-Hyb` reference model, the proposed `FBA-Hyb` framework demonstrates superior accuracy, increasing the average *R*^2^ over product (*P*) and biomass (*X*) by 19%. For benchmarking, the figure also includes results from `OptFed` [6], a purely mechanistic model specifically engineered for this process in a previous study. As `OptFed` was tailored to this specific process and dataset, it represents a strong benchmark that a hybrid model is not necessarily expected to surpass. Notably, the performance of `FBA-Hyb` is highly competitive with `OptFed`, exhibiting only marginal differences in predictive capability despite the hybrid nature of the former. Complementary to the reported *R*^2^ values, the combined NMAE for *P* and *X* are presented in Sup. Fig. S4. While the discrepancy in NMAE is less pronounced (0.22 for `Std-Hyb` vs. 0.20 for `FBA-Hyb`, i.e. 9%), `FBA-Hyb` still outperforms the reference model `Std-Hyb` (no values for `OptFed` model were reported in the original study).

**Figure 2:**
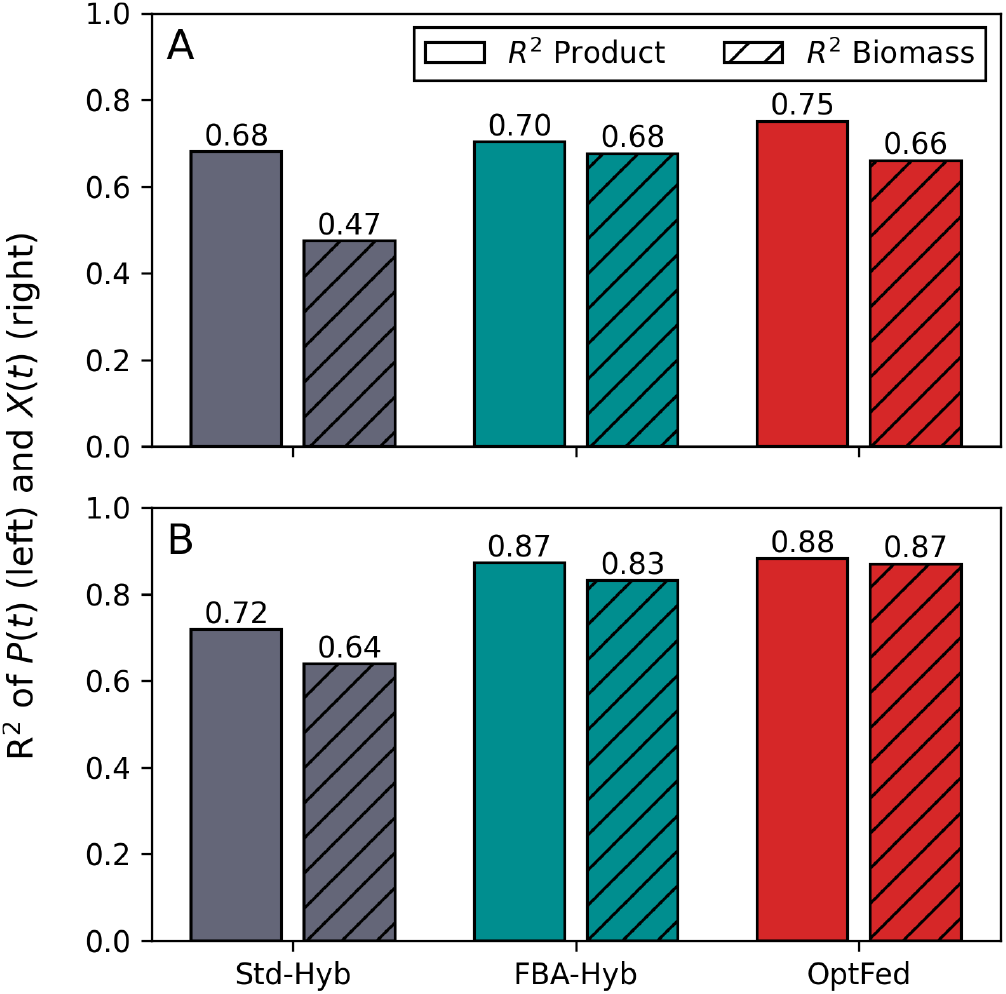
Validation results. **A** shows the *R*^2^ values for `Std-Hyb` and `FBA-Hyb` of the LOPO CV study while **B** shows their respective performance on an unseen test data set. For further comparison, `OptFed`, a mechanistic model developed by [6], is also included.

To test the models on unseen processes, we used three additional PROL fed-batch experiments that were not used for model building, hyperparameter selection, and cross validation. These test processes were run with different initial conditions, temperature and feeding profiles [6]. Predictions

In this test set, ⟨`FBA-Hyb`⟩_2_ again outperformed ⟨`Std-Hyb`⟩ by were made with the ensembles (indicated by ⟨⋅⟩, Sec. 2.8). an average increase in *R*^2^ of 20% and a decrease in NMAE of 44% (Fig. 2B and Sup. Fig. S3).

Furthermore, we tested the robustness of these results in a structural sensitivity analysis along two axes (Sup. Sec. S1.2 and Sup. Fig. S4): (i) the hybrid model architecture (`Std-Hyb`, `FBA-Hyb`, and `FBA-Hyb-SR`, in which ANN_1_ is additionally replaced by symbolic equations via SR), and (ii) the model configuration, i.e., how the base feed and the substrate uptake are modeled. In all configurations, `FBA-Hyb` was more accurate than `Std-Hyb` in the LOPO CV, with an average 42% increase in *R*^2^ and 21% reduction in NMAE. The configuration used throughout the main manuscript (`linear/none`) yielded the best `Std-Hyb` and the second-best `FBA-Hyb` variant. `FBA-Hyb-SR` performed slightly better than `FBA-Hyb` on average and had the best performing variant overall. However, as this gain was inconsistent across configurations, and `FBA-Hyb-SR` is considerably more computationally expensive, we focus on `FBA-Hyb` in the main manuscript (Sup. Sec. S1.2).

### 3.3. FBA-Hyb respects metabolic stoichiometry avoiding infeasible predictions

To illustrate why `FBA-Hyb` outperforms `Std-Hyb` in the LOPO CV, we sampled 1000 new data points out of the experimental distributions of the concentration (**c**) and control (**u**) for the PROL case study. From these sampled values we predicted metabolic fluxes (**q**) with the ensemble models ⟨`Std-Hyb`⟩ and ⟨`FBA-Hyb`⟩. The predicted values of **q**_*ℛ*_ were set as constraints of the original GSMM FBA model and the minimal substrate uptake requirement 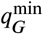 was calculated.

If the predicted 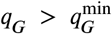 the mass balance was deemed violated and the ratio of 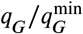 was used as the quantitative measurement of mass balance violation.

The results of this analysis are shown in Figure 3F. There, it is visible that over 22% of the `Std-Hyb` predictions of the sampled points resulted in infeasible GSMM FBA models. `Std-Hyb` violated metabolic network stoichiometry by an average quantity of 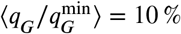 over all sampled points. Due to the architecture^*G*^ of the `FBA-Hyb` model that includes a very accurate surrogate FBA step there is virtually no infeasibility detected.

**Figure 3.**
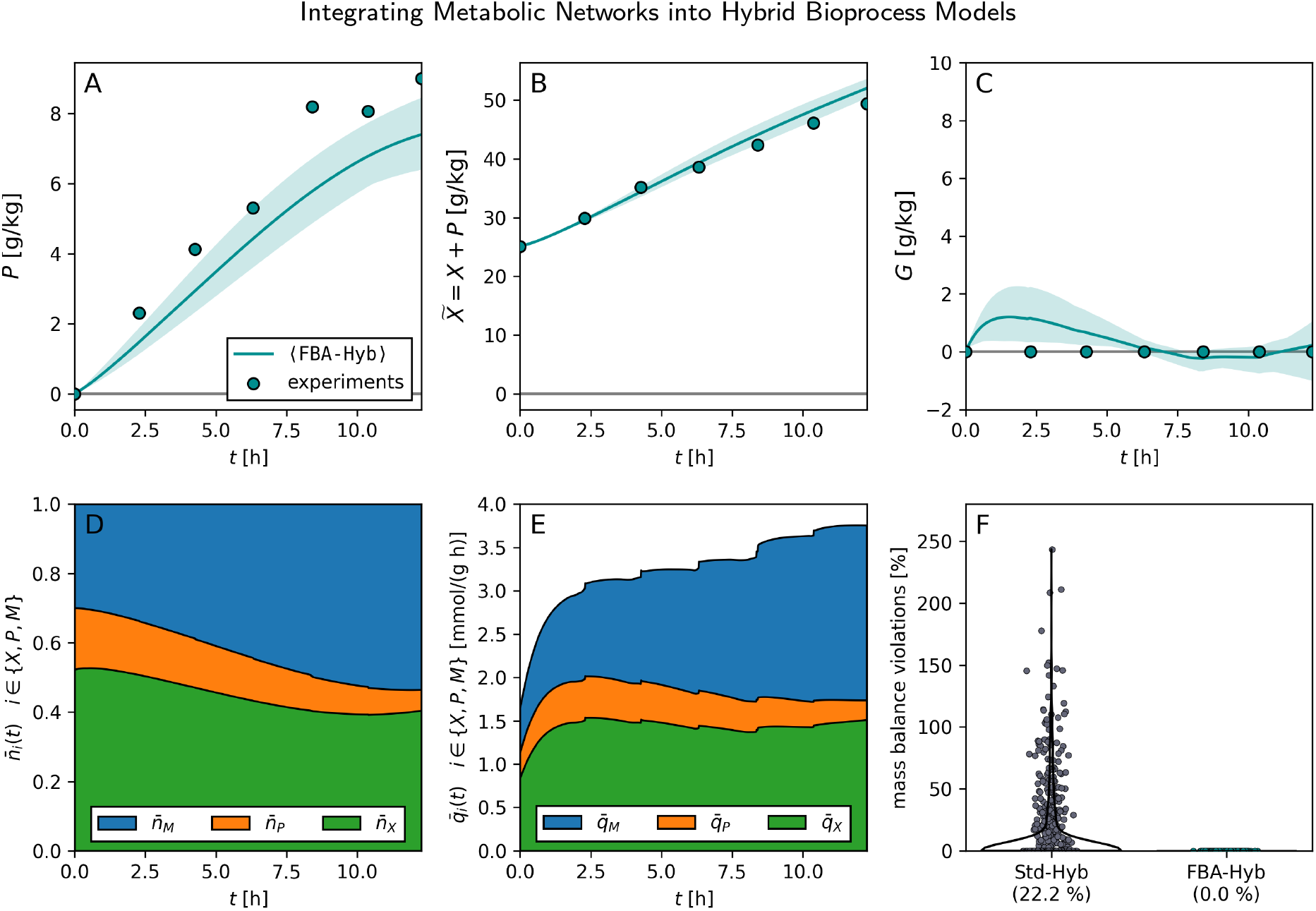
FBA-Hyb external state and internal rate predictions. **A, B, C** Measured (circles) and ⟨`FBA-Hyb`⟩ -predicted (lines) product, biomass, and substrate concentrations of one selected PROL process. Shaded areas indicate the minimum and maximum of the ensemble members. **D, E** Product-to-substrate normalized values of 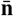 and 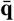, respectively, predicted by ⟨`FBA-Hyb`⟩ over time for the same process. Normalization as described in Sup. Sec. S1.3 was performed for easier interpretation. **F** Mass balance violations of ⟨`Std-Hyb`⟩ and ⟨`FBA-Hyb`⟩ for new data points sampled from the training data distribution.

### 3.4. FBA-Hyb predictions are stable

Fig. 3 shows the measured and predicted product, biomass, and glycerol substrate concentrations (panels A, B, C) as well as the predicted values of **n** (i.e., the FBA objective weights, panel D) and *q*_*i*_ (i.e., the metabolic fluxes of *i* ∈ {*X, P* , *M*}, panel E) over time. All values were predicted by the final ensemble ⟨`FBA-Hyb`⟩ for one selected PROL process in the training set. Product-to-substrate yield normalized values of **n** and **q** (i.e., 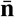, 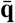) are shown to allow easier comparison (for details, see Sup. Sec. S1.3).

In this process, the product concentration is underestimated (*R*^2^ of 0.81, NMAE of 0.22), whereas the biomass concentration is predicted well (*R*^2^ of 0.96, NMAE of 0.03). The predicted glycerol concentration remains between −0.2 and 1.2 g kg^−1^, which is close to the measured value of zero, given that a total of 210 g of glycerol was fed.

In more detail, Fig. 3D illustrates the ensemble prediction output of the ⟨`FBA-Hyb`⟩ module ANN_1_ (**c u**; θ) (Eq. 6b). There it is visible that the proportions going towards biomass 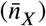 and product 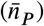 decrease over the process time by 23% and 66%, respectively, whereas the proportion going to the hydrolysis of ATP to ADP (i.e., the maintenance, 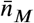) increases by 79%. Hence, an increasing share of the consumed substrate is predicted to be used for non-growth functions, for example, maintenance or stress responses due to the overexpression of recombinant proteins [42, 43]. Consequently, the biomass grows more slowly than the exponentially increasing feed, even though no glycerol accumulates (Fig. 3B and C).

Similar trends can be seen in Fig. 3E, where the metabolic flux rates 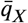 (green), 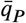 (orange), and 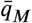 (blue) are plotted. There, also the total value of 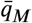 increases over time, whereas the production flux 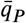 decreases towards the end of the process. However, there are discrete steps in the values over time for all three variables. These steps result from discrete volume changes that happen due to sampling of the bioreactor and have to be expected when working with carbon-limited processes (i.e., the glycerol concentration close to zero in Fig. 3C). Similar steps can be seen in the prediction of **q** from ⟨`Std-Hyb`⟩ (Fig. S5). Although the predictions look similar, direct predictions of discrete steps with ANN may become problematic during training [44]. The ⟨`FBA-Hyb`⟩ architecture avoids this direct prediction of discrete steps completely, resulting in smooth predictions of 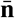 (Fig. 3D).

### 3.5. FBA-Hyb models nutrient limitations

As a second case study, we considered the pDNA production process under sulfate limitation [34]. In this system, sulfate depletion was shown to act as a key growth-limiting nutrient and to decouple product formation from biomass growth [34]. Here we tested the performance of `Std-Hyb` and `FBA-Hyb` on predicting this behavior.

Importantly, sulfate concentrations were not measured over time in the experiments, only initial concentrations are available. Therefore, sulfate uptake rates cannot be trained directly. Within the `Std-Hyb` framework, incorporating such an unobserved nutrient is not straightforward, since it would require either additional measurements or an explicit mechanistic balance equation for sulfate. In contrast, the `FBA-Hyb` formulation naturally accommodates this situation. Because sulfate uptake is already represented by a flux *q*_*S*_ ∈ **q**_*ε*_, it can be included in the flux balance constraint set **q** as long as the initial medium concentration is known, which is the case here. This allows sulfate availability to implicitly constrain growth and production through the learned fluxes without requiring time-resolved sulfate measurements.

Fig. 4 summarizes the resulting model behavior. Panels A-C show the ensemble ⟨`FBA-Hyb`⟩ predictions (lines) and experimental measurements (circles) for product (*P*), biomass (*X*), and sulfate (*S*) concentrations in the bioreactor, respectively. The different colors correspond to initial SO^2−^concentrations of 12, 16, 20, and 29 mmol L^−1^. Experim^4^ ental training data was available for *S*(0) = 16 and 29 mmol L^−1^. The vertical dotted lines mark the time points of maximal biomass for each condition. Panels D and E of Fig. 4 show the predicted values of the FBA objective weights (**n**) and the metabolic fluxes (**q**) for a selected sulfate-limiting process (*S*(0) = 16 mmol L^−1^). By comparing panels C and D it is visible that `FBA-Hyb` reduces the FBA objective weight for biomass (*n*_*X*_), and subsequently *q*_*X*_, once sulfate becomes limiting. This is consistent with sulfur being required for biomass but not for pDNA synthesis [34]. ⟨`FBA-Hyb`⟩ correctly captures the experimentally observed reduction in biomass growth as sulfate becomes limiting, together with the continued increase in product concentration.

**Figure 4:**
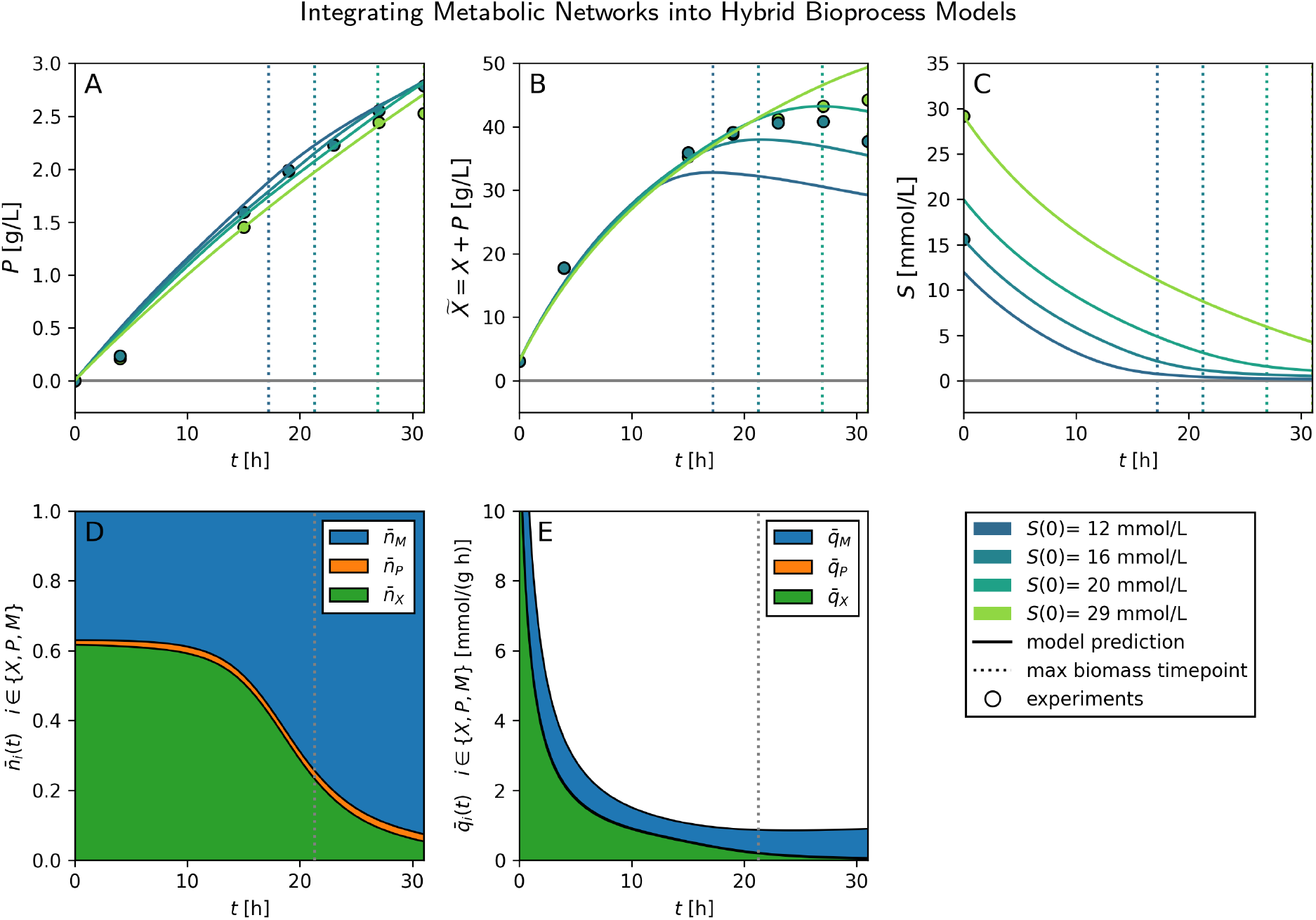
Sulfate limitation during process optimization. Panels **A, B, C** show predicted state variables by `FBA-Hyb` for different initial concentrations of SO^2−^. Panels **D** and **E** illustrate ⟨`FBA-Hyb`⟩‘s predicted normalized FBA objective weights 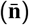 and metabolic fluxes 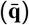, respectively. Note that in panel **E** the orange area is too small to be visible.

For comparison, Sup. Fig. S6A and B show the corresponding predictions of *X* and *P* obtained with the ⟨`Std-Hyb`⟩ model. There, the initial concentration of sulfate (*S*(0)) is part of the ANN input via the control vector **u** (Eq. 11d), however time-dependent concentrations of sulfate cannot be tracked. ⟨`Std-Hyb`⟩ reproduces the sulfate limitation from the training data reasonably well, but when reducing *S*(0) even further, it fails. In panel C of Sup. Fig. S6, we visualized this by post-hoc calculating the hypothetical *S*(*t*) consumption with the original GSMM FBA model which shows that for *S*(0) = 12 mmol L^−1^ ⟨`Std-Hyb`⟩ predicts biomass concentrations that would require more sulfate than is available in the medium. Because sulfate is not modeled dynamically, ⟨`Std-Hyb`⟩ was unable to learn the correct limiting behavior, leading to physiologically infeasible trajectories.

Overall, this case study demonstrates that `FBA-Hyb` can learn nutrient-limited growth behavior directly from experimental data and initial medium compositions, even when the limiting substrate is not measured over time. A key strength of the `FBA-Hyb` framework is that sulfate uptake fluxes are embedded in **q**_*ℛ*_ and predicted through the surrogate FBA, allowing the model to dynamically adjust the underlying metabolic flux distribution. Importantly, the hybrid model does not merely interpolate the observed data, but instead, adapts the objective weights in the surrogate FBA input in response to changing nutrient availability. This enables `FBA-Hyb` to capture the decoupling of biomass growth and product formation under sulfate limitation in a consistent manner. By learning these biologically meaningful patterns, the model can generalize beyond the training conditions and predict growth and production dynamics for sulfate concentrations that were not measured. This highlights the potential of `FBA-Hyb` to describe such metabolic shifts directly from process data while respecting the network stoichiometry, offering both predictive power and insight into cellular behavior under nutrient-limited conditions.

## 4. Discussion

### 4.1.The FBA surrogate model

The approximation of FBA solutions via symbolic regression is remarkably robust, as it uncovers a generalized equation (Eq. 12) that is structurally analogous to the yield coefficients traditionally employed in bioprocess modeling. The physical interpretability of this formulation can be exemplified by setting the objective weight vector to **n** = (1, 0, 0)^T^, which reduces the generalized surrogate model to:

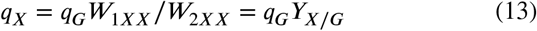

where *Y*_*X*/*G*_ represents the true substrate-to-biomass yield of the cell [9]. In contrast, while previous studies have utilized FBA surrogate models, these have predominantly relied on ANN-based architectures [29, 23]. While functional, blackbox approaches do not yield the physically meaningful and easy-to-evaluate mathematical expressions provided by SR.

### 4.2. The FBA-Hyb architecture

The dynamic modification of the FBA objective function in this study is inspired by findings that cellular objectives are often state-specific and vary according to the environmental context [45]. This observation aligns with previous efforts to predict FBA objectives to improve model accuracy [46]. However, in `FBA-Hyb`, **n** is not interpreted as an intrinsic objective of the cell, but as a parameterization of the partitioning of the carbon substrate among biomass, product, and ATP hydrolysis on the set of stoichiometrically optimal flux distributions [47]. Thereby, the ATP maintenance fluxacts as a lumped sink for substrate that is respired without contributing to biomass or product formation, e.g., due to non-growth-associated maintenance or stress responses [42]. Changes in **n** should therefore be regarded as a phenomenological proxy for unmodeled regulatory effects (e.g., proteome allocation or metabolic burden), rather than as evidence that cells actively adapt their metabolic objective. A primary advantage of the `FBA-Hyb` approach compared similar methodologies is the structural integration of FBA within the hybrid model. This ensures that mass balances are explicitly maintained during both parameter training and subsequent process optimization, even when extrapolating beyond what was explicitly trained. In contrast, methods that validate FBA predictions only post-training may lack such inherent consistency [25]. This challenge is also observed in PINNs, where physical relationships are learned as soft constraints; if predictions deviate significantly from the training data, the model may yield nonphysical results and suffer from decreased performance [21]. In many FBA-base bioprocess models (e.g., dynamic

In many FBA-base bioprocess models (e.g., dynamic FBA) overflow metabolism and production rates are implemented as constraints rather than being part of the objective function [34]. However, this can lead to infeasibilities that have to be handled cautiously [13]. The architecture of `FBA-Hyb` ensures that infeasibilities within the metabolic network cannot arise during simulation. This provides a computational advantage over frameworks such as NEXT-FBA, which may require solving a mixed-integer linear program (MILP) to relax ANN predictions when they lead to metabolic infeasibility [24]. By maintaining feasibility by design, `FBA-Hyb` avoids the need for additional optimization-based relaxation steps.

### 4.3. The prediction of n

The results presented in Fig. 3D and E highlight the advantages of predicting the FBA objective weights (**n**) rather than the metabolic rates (**q**) directly. In carbon-limited processes, discrete changes in the substrate uptake rate, e.g., due to feed or volume changes, are directly reflected in **q** (Fig. 3E). In `FBA-Hyb`, these changes are propagated through the surrogate FBA, such that **n** itself can remain smooth (Fig. 3D). In the context of hybrid modeling, ANNs trained to predict kinetic rates (e.g., `Std-Hyb`) often require high sensitivity to input variables to capture these dynamics, which can lead to extreme or non-physical predictions when training data are sparse (a common challenge in bioprocess development). Conversely, predicting **n** provides a more stable target for the ANN, as it changes more smoothly over time, allowing for more effective regularization and robust generalization [44]. As objective functions inferred from flux data are generally not unique [48], a physiological validation of the predicted **n** would require intracellular data such as ^13^C-based metabolic flux analysis or proteomics [47], which is beyond the scope of this study.

Under this framework, only the primary substrate uptake rate, *q*_*G*_, must be predicted directly. This simplification is justified by the fact that often follows predictable patterns: it typically operates near maximum velocity in the presence of excess substrate due to low Michaelis-Menten constants [49], or is governed by the feed rate in carbon substrate-limited conditions, such as in the SLIM case study (Eq. 11f) [34]. While these trends may deviate during overflow metabolism, such regimes are generally avoided in optimized industrial bioprocesses, making *q*_*G*_ a relatively straightforward feature for the model to learn.

### 4.4. Comparison to other methods

dFBA is a well-established method for bioprocess modeling. However, historically, the objective reaction has been treated as static, i.e., a single value for **n** is assumed over the entire process duration [12]. While this approximation may be adequate for some processes, it does not hold in our case. As shown in Sup. Fig. S7, we evaluated the effect of fixing **n** in `FBA-Hyb` to a constant vector, instead of predicting it dynamically using an ANN, on LOPO cross-validation performance. In both case studies, this modification led to a deterioration in *R*^2^ and NMAE, reducing predictive accuracy even below that of `Std-Hyb`. These results highlight the synergistic advantage of the `FBA-Hyb` architecture, which outperforms both conventional hybrid models (`Std-Hyb`) and dFBA-like approaches with static **n**.

In terms of LOPO CV performance, `FBA-Hyb` and `OptFed` demonstrate comparable predictive accuracy for the PROL case (Fig. 2). This is expected, as a well-validated mechanistic model that captures all relevant process effects is generally not outperformed by a hybrid model. The advantage of hybrid models rather lies in processes where mechanistic descriptions fail to capture all observed effects [16]. However, `OptFed` is a highly specialized, purely mechanistic model that is specifically tailored to the process configuration and control structure considered here. Consequently, extending `OptFed` to include additional control variables would require substantial model reformulation [6]. In contrast, the `FBA-Hyb` framework is inherently more flexible; additional control inputs, such as pH and dissolved oxygen, can be incorporated by adding them to the input of the data-driven components (i.e., ANN_1_ and ANN_2_) without modifying the underlying constraint structure. This modularity represents a key practical advantage of `FBA-Hyb` over purely mechanistic approaches while maintaining similar predictive performance.

While Bogaerts et al. previously proposed a method to predict FBA objectives dynamically, there are significant differences compared to the present approach [27]. Their methodology is entirely mechanistic and does not utilize machine learning, which can make the search for an appropriate model structure more difficult and time-intensive. This is particularly relevant when including non-standard control variables that cannot be easily described by traditional enzyme kinetic-like terms.

Our `FBA-Hyb` offers a distinct structural advantage during model training over sequential hybrid frameworks, such as the digital-twin approach described by Richelle et al. [26]. In that framework, the authors chose to couple ANNs, kinetic ODEs, and a constraint-based FBA module in a nondifferentiable manner, with the FBA component acting as an external solver. This prevents the propagation of a learning signal to the upstream ANNs during training. In contrast, `FBA-Hyb` integrates the FBA module more tightly into the learning process. By employing a differentiable surrogate for the inner FBA problem, backpropagation is enabled through the entire model, allowing for the end-to-end optimization of the hybrid architecture. Only with this architecture, `FBA-Hyb` is able to accurately learn metabolic shifts from unmeasured metabolites (Fig. 4D).

### 4.5. Outlook

The flexibility of the `FBA-Hyb` framework allows for systematic control over the degree of hybridization [50]. Models with a high degree of hybridization have a bigger emphasis on ANNs, more flexibility, more parameters and consequently need more data to train. Models with low degree of hybridization emphasize mechanistic equations resulting in the opposite behavior [50]. In the context of `FBA-Hyb`, by adjusting the composition of the objective weight vector, **n**, used to train the surrogate FBA model the degree of hybridization can be adjusted. When prior knowledge suggests that specific fluxes, such as the maintenance rate *q*_*M*_ , remain approximately constant or not enough training data is available, they can be excluded from the objective vector (e.g., **n**^′^ = (*n*_*X*_, *n*_*P*_)^T^). This approach increases the proportion of explicitly encoded process knowledge through the surrogate FBA, thereby constraining the prediction space of the datadriven components and potentially enhancing robustness by mitigating overfitting. Conversely, the model can be shifted toward a more data-driven formulation by expanding the objective vector to include additional fluxes, such as byproduct synthesis (e.g., acetate *A*, **n**^′′^ = (*n*_*X*_, *n*_*P*_ , *n*_*M*_ , *n*_*A*_)^T^). While this expansion increases model expressivity and reduces reliance on prior mechanistic assumptions, it also requires more bioprocess data for robust model training. Moreover, increasing the size of **n** increases the combinatorial complexity when generating the training data required for the surrogate FBA model.

In the current implementation of `FBA-Hyb`, the surrogate model is primarily utilized to predict exchange fluxes (Eq. 12). However, this methodology can be extended to estimate internal metabolic fluxes as well. Incorporating information regarding these internal fluxes from previous timesteps as inputs to ANN_1_ (Eq. 6c) could provide the network with more granular state information, potentially improving predictive performance in future iterations.

A current limitation of the `FBA-Hyb` framework is the handling of certain hard constraints, such as the positivity of chemical concentrations (*S* ≥ 0). As illustrated in Fig. 4C, minor violations of these constraints may occur because they are currently implemented as soft constraints via penalty terms in the loss function, which can be challenging to balance during training. Reformulating `FBA-Hyb` as a system of differential-algebraic equations (differential algebraic equation (DAE)) could facilitate the more rigorous integration of such constraints [51]. However, such a reformulation is beyond the scope of the present study.

## 5. Conclusion

This study introduces `FBA-Hyb`, a novel hybrid modeling framework that tightly integrates metabolic networks into a fully differentiable architecture. By employing a symbolic surrogate of the FBA problem, the framework ensures strict adherence to mass balances while allowing easy backpropagation and maintaining the flexibility of data-driven approaches. Our results demonstrate that `FBA-Hyb` outperforms standard hybrid models (`Std-Hyb`) by providing more stable predictions, even when training data are sparse, and by allowing for the seamless inclusion of untracked chemical species as state variables. The modularity of the `FBA-Hyb` framework, coupled with its ability to maintain metabolic feasibility by design, offers a robust alternative to existing hybrid methods.

## Author contribution

**Mathias Gotsmy**: Conceptualization, Data curation, Formal analysis, Investigation, Methodology, Software, Validation, Visualization, Writing – original draft. **Gonzalo Guillén Gosálbez**: Conceptualization, Funding acquisition, Methodology, Project administration, Resources, Supervision, Writing – review and editing.

## Declaration of competing interests

The authors do not declare any competing interests.

### Notation list

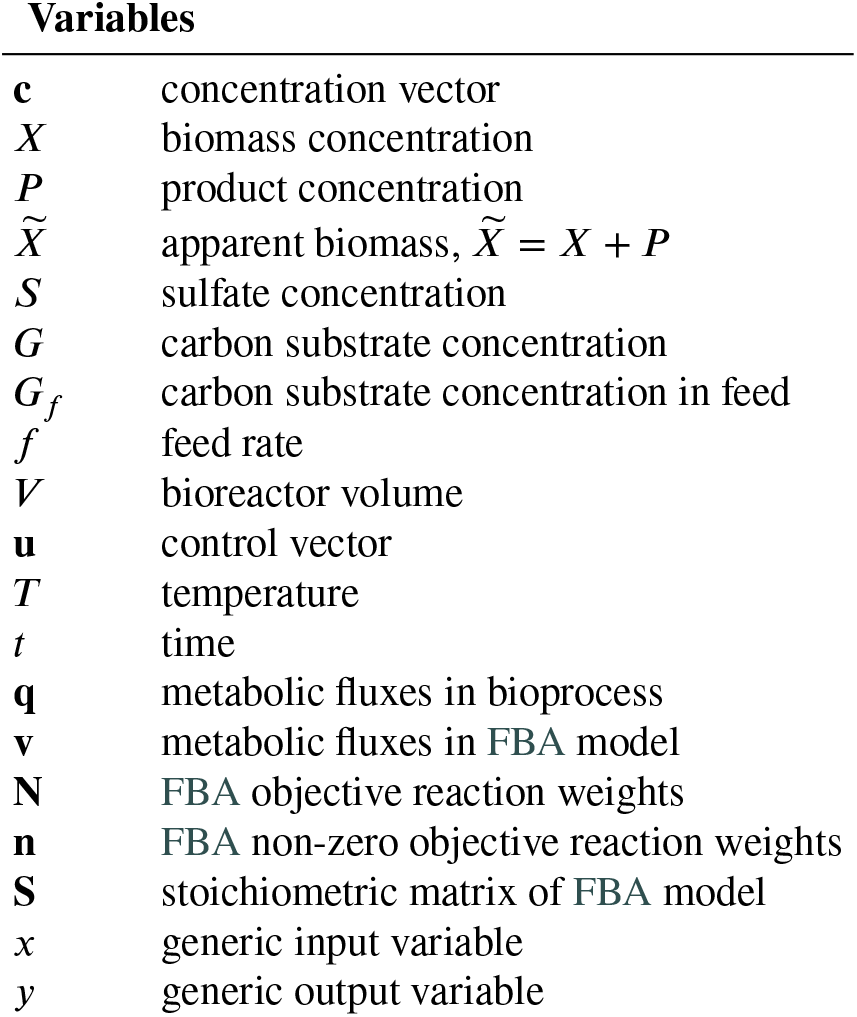
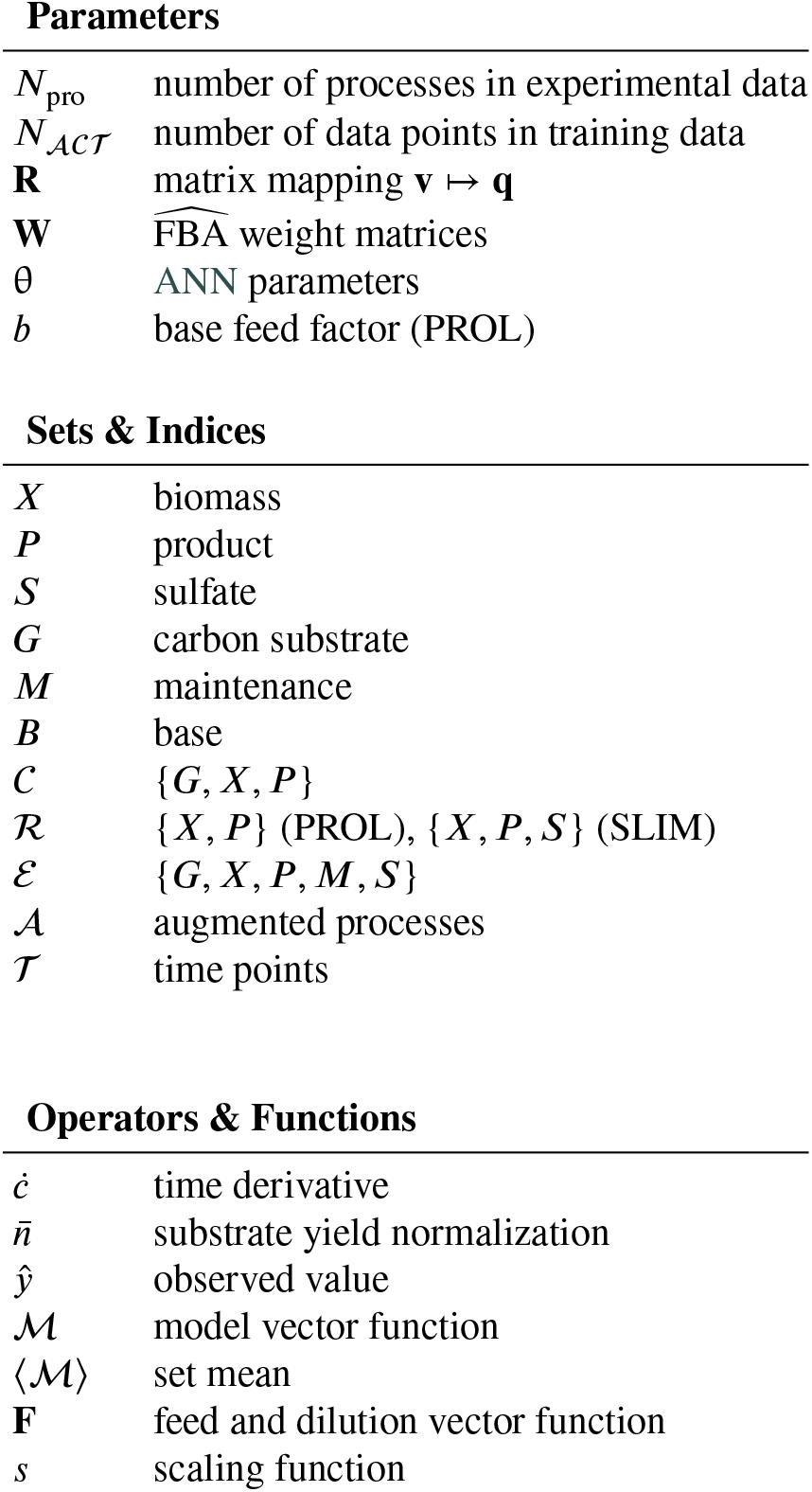

| Variables |  |
| --- | --- |
| $\mathbf{c}$ | concentration vector |
| $X$ | biomass concentration |
| $P$ | product concentration |
| $\tilde{X}$ | apparent biomass, $\tilde{X} = X + P$ |
| $S$ | sulfate concentration |
| $G$ | carbon substrate concentration |
| $G_f$ | carbon substrate concentration in feed |
| $f$ | feed rate |
| $V$ | bioreactor volume |
| $\mathbf{u}$ | control vector |
| $T$ | temperature |
| $t$ | time |
| $\mathbf{q}$ | metabolic fluxes in bioprocess |
| $\mathbf{v}$ | metabolic fluxes in FBA model |
| $\mathbf{N}$ | FBA objective reaction weights |
| $\mathbf{n}$ | FBA non-zero objective reaction weights |
| $\mathbf{S}$ | stoichiometric matrix of FBA model |
| $x$ | generic input variable |
| $y$ | generic output variable |
| Parameters |  |
| $N_{\text{pro}}$ | number of processes in experimental data |
| $N_{\text{ACT}}$ | number of data points in training data |
| $\mathbf{R}$ | matrix mapping $\mathbf{v} \mapsto \mathbf{q}$ |
| $\mathbf{W}$ | FBA weight matrices |
| $\theta$ | ANN parameters |
| $b$ | base feed factor (PROL) |
| Sets & Indices |  |
| $X$ | biomass |
| $P$ | product |
| $S$ | sulfate |
| $G$ | carbon substrate |
| $M$ | maintenance |
| $B$ | base |
| $C$ | $\{G, X, P\}$ |
| $\mathcal{R}$ | $\{X, P\}$ (PROL), $\{X, P, S\}$ (SLIM) |
| $\mathcal{E}$ | $\{G, X, P, M, S\}$ |
| $\mathcal{A}$ | augmented processes |
| $\mathcal{T}$ | time points |
| Operators & Functions |  |
| $\dot{\phantom{x}}$ | time derivative |
| $\bar{n}$ | substrate yield normalization |
| $\hat{y}$ | observed value |
| $\mathcal{M}$ | model vector function |
| $\langle \mathcal{M} \rangle$ | set mean |
| $\mathbf{F}$ | feed and dilution vector function |
| $s$ | scaling function |

## Notes

https://github.com/Gotsmy/FBA-Hyb

### Competing Interest Statement

The authors have declared no competing interest.

### Summary of Updates

Revised after peer review: added a hold-out validation on three unseen fed-batch processes, clarified that the FBA objective weights are a data-driven approximation rather than a physiological cellular objective, extended Figure 3 with measured concentrations, and clarified the performance reporting and architecture comparison.

