## Supplemental Material for "Integrating Metabolic Networks into Hybrid Bioprocess Models"

### S1. Supplementary Notes

#### S1.1. Bioreactor Volume Normalization

Discrete jumps in the bioreactor volume can happen due to sampling (i.e., reduction of the sampling volume) or feeding (i.e., bolus feeds). Normalization can be performed to move this discrete jumps from the ordinary differential equation (ODE) state variables to the control variables, which is typically easier to handle for numerical solvers. Here, a method developed by [6] is described.

Given an ordered sequence of timepoints $\mathcal{S} = (t_1, \dots, t_n)$, where discrete volume removals $\mathcal{V} = (V_{S1}, \dots, V_{Sn})$ occur, with $V_{Si} > 0$, and a non-normalized reactor volume $\widehat{V}$ the normalized reactor volume is defined by:

$$V(t) = \frac{\widehat{V}(t)}{s(t)} \tag{S1a}$$

where the normalization factor is

$$s(t) = \prod_{\substack{1 \le i \le n \\ t_i < t}} \left(1 - \frac{V_{Si}}{\widehat{V}(t_i^-)}\right) \tag{S1b}$$

with $s(t) = 1$ if $t \le t_1$. Analogously, any feed rate $\widehat{f}$ transforms as

$$f(t) = \frac{\widehat{f}(t)}{s(t)}. \tag{S1c}$$

#### S1.2. Hybrid Model Architecture and Configuration Comparison

To test the robustness of our results, 18 model variants were evaluated for the PROL case study along two axes: (i) the hybrid model architecture (`Std-Hyb`, `FBA-Hyb`, and `FBA-Hyb-SR`), and (ii) the model configuration, i.e., the combination of a base feed model (`linear` or `ann`) and a substrate uptake constraint (`none`, `sigmoid`, or `min`). Configurations are denoted as `base feed/uptake constraint`, e.g., `linear/none`.

##### *S1.2.1. Methods*

*Configuration: Base Feed Model* Here, we implemented two methods of predicting the base feed ($f_B$) as defined in Eq. 11a. The first method is a linear model as suggested by [6], i.e.,

$$f_B = b\, q_X \qquad \texttt{(linear)} \tag{S2a}$$

and the second version is an artificial neural network (ANN)

$$f_B = \mathrm{ANN}(\mathbf{c}, \mathbf{u}, q_X; \theta) \qquad \texttt{(ann)} \tag{S2b}$$

where parameters $b$ and $\theta$ are estimated during training.

*Configuration: Substrate Uptake Constraint* In both, `Std-Hyb` and `FBA-Hyb`, an ANN predicts a value of $q_G$ dynamically as

$$q_G^{\mathrm{ANN}} = \mathrm{ANN}(\mathbf{c}, \mathbf{u}; \theta) \tag{S3a}$$

however, if the substrate concentration $G$ is low, the maximum $q_G$ is governed by the substrate feed rate ($f_G G_f$) and the total bioreactor biomass ($XV$) as

$$q_G^{\max} = f_G G_f / (XV) \qquad \text{if } G \to 0. \tag{S3b}$$

To test the influence of adding this information to the model prediction, we developed three methods

$$q_G = q_G^{\mathrm{ANN}} \qquad \texttt{(none)} \tag{S3c}$$

$$q_G = \alpha q_G^{\mathrm{ANN}} + (1-\alpha) q_G^{\max} \quad \text{with } \alpha = \mathrm{sigmoid}((G-\theta_1)/\theta_2) \qquad \texttt{(sigmoid)} \tag{S3d}$$

$$q_G = \min(q_G^{\mathrm{ANN}}, q_G^{\max}) \qquad \texttt{(min)} \tag{S3e}$$

where parameters $\theta_1$ and $\theta_2$ are estimated during training.

*Architecture:* FBA-Hyb-SR Symbolic regression (SR) can be advantageous over black box models (like ANNs) due to better interpretability and generalization [30]. Here, SR is used to transform the $\mathrm{ANN}_1$ to symbolic equations (indicated as SRE) resulting in a adapted form of the FBA-Hyb model:

$$\dot{\mathbf{c}}(t) = \mathbf{q}X + f/V(\mathbf{c}_f - \mathbf{c}) \tag{S4a}$$
$$\mathbf{n} = \mathrm{SRE}(\mathbf{c}, \mathbf{u}; \theta_3) \qquad \text{(FBA-Hyb-SR)} \tag{S4b}$$
$$q_G = \mathrm{ANN}_2(\mathbf{c}, \mathbf{u}; \theta_2) \tag{S4c}$$
$$\mathbf{q}^\mathrm{T} = (q_G,\ \mathbf{q}_\mathcal{R}^\mathrm{T}) \tag{S4d}$$
$$\mathbf{q}_\mathcal{R} = \mathbf{R}\ \widehat{\mathrm{FBA}}(q_G, \mathbf{n}), \tag{S4e}$$

referred to as FBA-Hyb-SR. Training data for SR were obtained by tracking the inputs and outputs of $\mathrm{ANN}_1$. These data, collected across all bioprocess training simulations, were then used to fit the symbolic equations, i.e., SRE (Eq. S4b). To maintain structural simplicity and physical interpretability, the symbolic regression search was restricted to $\{+, \times, \div\}$ operators with structural constraints implemented to prohibit nested divisions. These nesting rules prioritize the discovery of rational functional forms by allowing polynomial terms in numerators and denominators while preventing the evolution of overly complex recursive structures.

##### *S1.2.2. Results*

A leave-one-process-out (LOPO) cross-validation (CV) analysis as described in the main manuscript (Sec. 2.7) was performed for all 18 variants (Fig. S4). The results show that the linear/none configuration performed very well for both, Std-Hyb and FBA-Hyb, in terms of $R^2$ and normalized mean absolute error (NMAE) (Fig. S4). Therefore, all further investigations were performed with this configuration. Going from Std-Hyb to FBA-Hyb improved the model performance in all configurations. The average improvements were 41.9 % in $R^2$ and −21.1 % in NMAE. Furthermore, going from FBA-Hyb to FBA-Hyb-SR lead to an average improvement of 2.5 % in $R^2$ and a decline of 0.014 % in NMAE.

##### *S1.2.3. Discussion*

Among the evaluated configurations, linear/none yielded the top-performing Std-Hyb and FBA-Hyb-SR models and the second-highest accuracy for the FBA-Hyb framework. These results underscore the predictive capacity of the ANN component, which, in the PROL case study, captured $q_G$ dynamics more effectively than alternative physics-driven implementations. While the FBA-Hyb-SR model in the linear/none configuration achieved the highest overall performance, the marginal average gains relative to FBA-Hyb did not justify the substantial increase in computational overhead and methodological complexity associated with symbolic regression. Moreover, the effect of the SR step was inconsistent across configurations, ranging from −1.8 % to +7.6 % in $R^2$. To prioritize model parsimony and mitigate the risk of overfitting the CV results, we chose to focus the main manuscript on the Std-Hyb and FBA-Hyb architectures in the linear/none configuration (in the main manuscript just referred to as Std-Hyb and FBA-Hyb). These selected variants are distinguished by a hatched fill in Fig. S4 (Fig. S4).

#### S1.3. Normalization of n and q

As they are proportional to the molar mass of the reaction product(s) (e.g., biomass, bioprocess product, maintenance) in a metabolic pathway of a genome-scale metabolic model (GSMM), **n** and **q** may take different orders of magnitudes. This makes direct comparison hard, therefore, for analysis, we normalized their values by their respective product-to-substrate yield $Y_{i/G}$,

$$\bar{n}_i = \frac{n_i Y_{i/G}^{-1}}{\sum_j \bar{n}_j} \tag{S5a}$$
$$\bar{q}_j = q_j Y_{j/G}^{-1} \tag{S5b}$$
$$Y_{k/G} = q_k / q_G \qquad i, j, k \in \{X, P, M\}. \tag{S5c}$$

The normalization is indicated by a bar and $\bar{\mathbf{n}}$ is again normalized to 1. The product-to-substrate yield $Y_{i/G}$ was calculated from the original GSMM from the PROL and SLIM case studies, respectively.

#### S2. Supplementary Tables

**Table S1:** Fitted parameters of $\mathbf{W}_1$ and $\mathbf{W}_2$ the PROL case study flux balance analysis (FBA) surrogate model (Eq. 12).

| $\mathbf{W}_1$ | | | $\mathbf{W}_2$ | | |
|---|---|---|---|---|---|
| 0.7393 | 0.2980 | 0.3161 | 0.7393 | 0.2980 | 0.3161 |
| 0.0427 | 0.0000 | 0.0000 | 0.7393 | 0.2980 | 0.3161 |
| 0.0000 | 0.0004 | 0.0000 | 0.7393 | 0.2980 | 0.3161 |
| 0.0000 | 0.0000 | 4.2678 | 0.7393 | 0.2980 | 0.3161 |
| 0.0276 | 0.0011 | 0.0000 | 1.8962 | 0.7635 | 0.8107 |

**Table S2:** Fitted parameters of $\mathbf{Y}$ and $\mathbf{W}$ the SLIM case study FBA surrogate model (Eq. 12).

| $\mathbf{Y}$ | | | $\mathbf{W}$ | | |
|---|---|---|---|---|---|
| 1.1091 | 0.0628 | 0.8564 | 1.1091 | 0.0628 | 0.8564 |
| 0.1006 | 0.0000 | 0.0000 | 1.1091 | 0.0628 | 0.8564 |
| 0.0000 | 0.0020 | 0.0000 | 1.1091 | 0.0628 | 0.8564 |
| 0.0000 | 0.0000 | 20.1250 | 1.1091 | 0.0628 | 0.8564 |
| 0.0710 | 0.0000 | -0.0000 | 3.1119 | 0.1769 | 2.3969 |

#### S3. Supplementary Figures

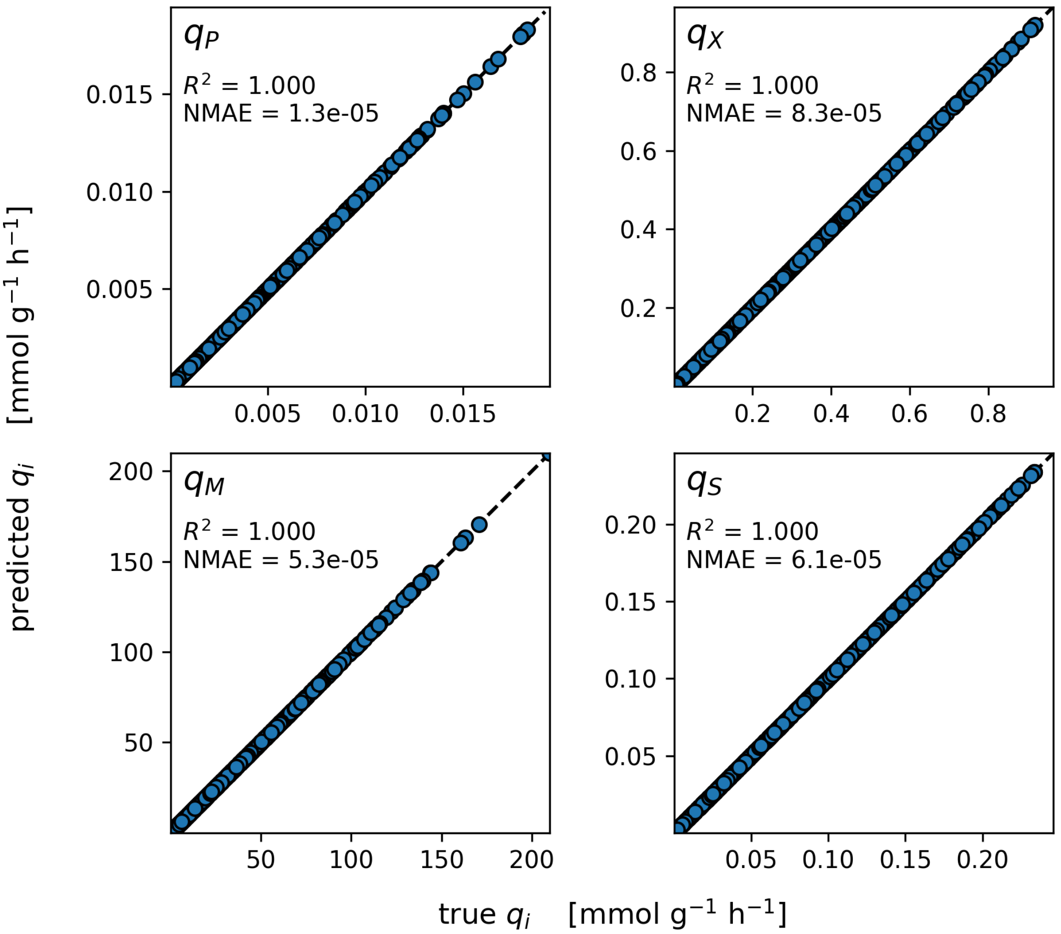


**Figure S1:** Performance of the surrogate FBA model on validation data for the PROL case study. It is clear that the recovered Eq. 12 and the fitted parameters reflect the FBA solution well and the surrogate modeling error is negligible. The plot of $q_G$ was omitted as the equation reduces to $q_G = (1, 0, 0, 0, 0)\widehat{\mathrm{FBA}}(q_G, \mathbf{n}) = q_G$ with machine precision.

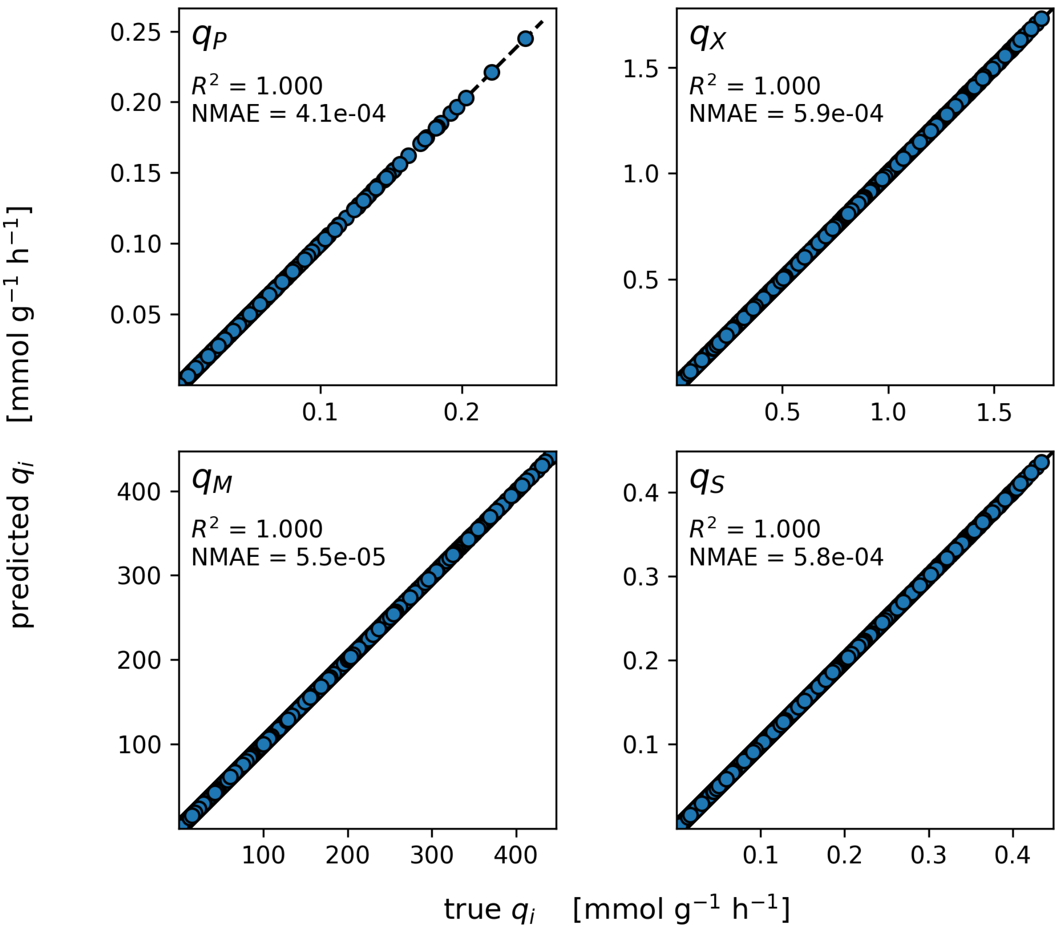


**Figure S2:** Performance of the surrogate FBA model on validation data for the SLIM case study. It is clear that the recovered Eq. 12 and the fitted parameters reflect the FBA solution well and the surrogate modeling error is negligible. The plot of $q_G$ was omitted as the equation reduces to $q_G = (1, 0, 0, 0, 0)\widehat{\text{FBA}}(q_G, \mathbf{n}) = q_G$ with machine precision.

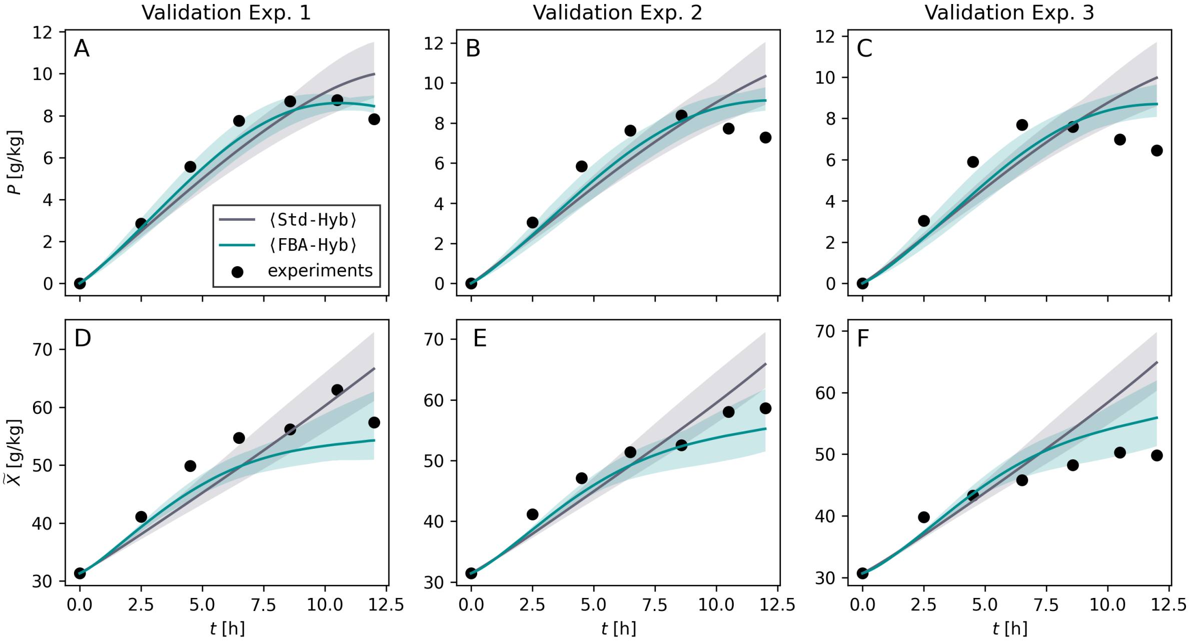


**Figure S3: Test set predictions.** Validation of Std-Hyb and FBA-Hyb on unseen PROL test data. The lines represent the LOPO CV ensemble mean and the shaded area the ensemble minimum and maximum. Performance metrics were calculated for the mean values.

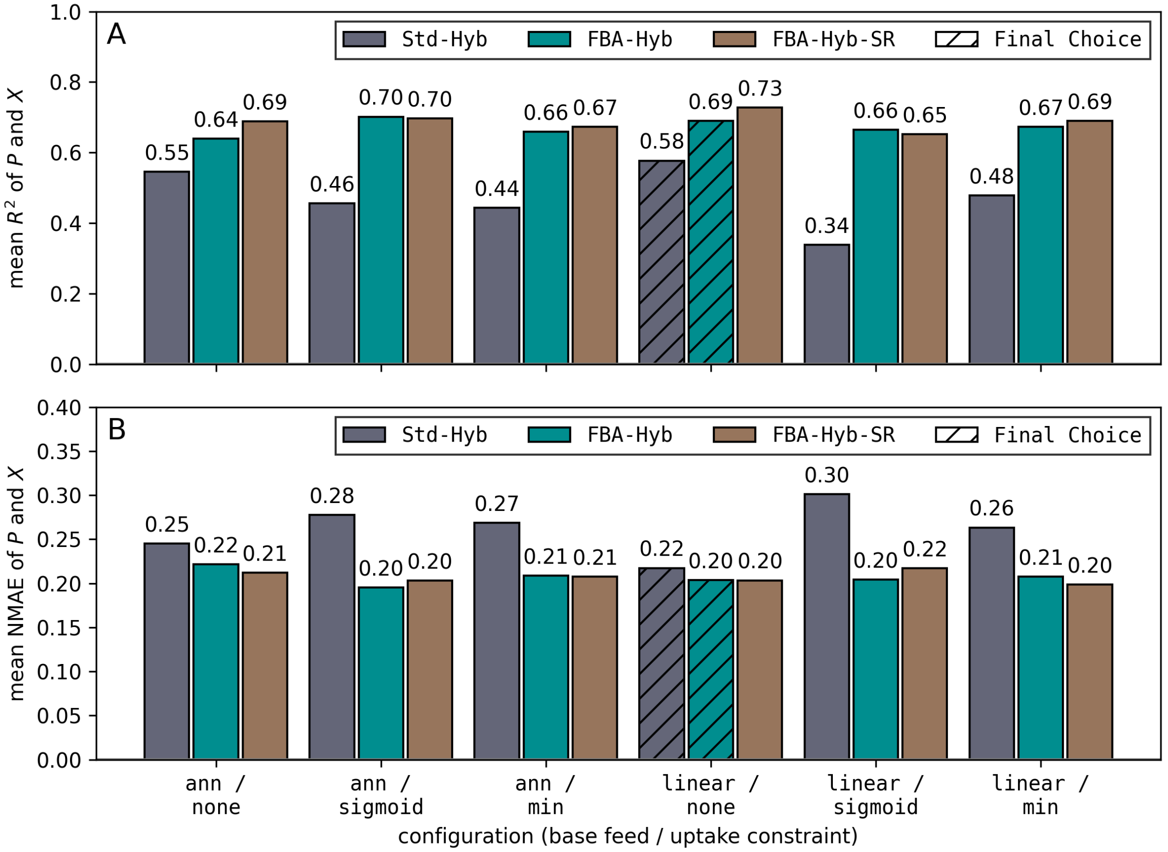


**Figure S4: Model architecture and configuration comparison.** LOPO CV $R^2$ (**A**) and NMAE (**B**) of the three hybrid model architectures (colors) in six configurations (x-axis; base feed model/substrate uptake constraint), as described in Sup. Sec. S1.2. Hatched bars mark the `Std-Hyb` and `FBA-Hyb` variants used in the main manuscript.

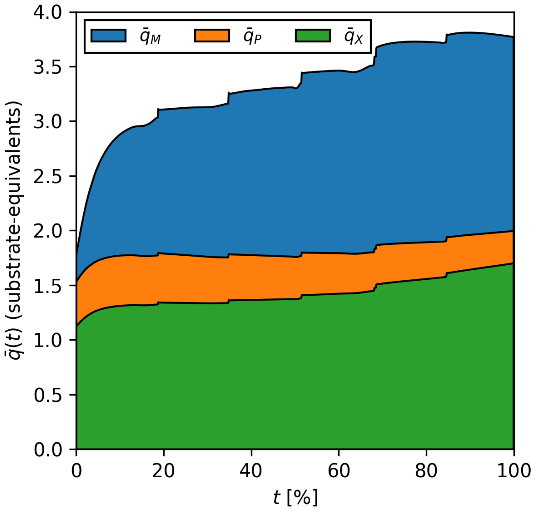


**Figure S5: `Std-Hyb` nternal rate predictions.** Product-to-substrate normalized values of $\bar{\mathbf{q}}$ predicted by ⟨`Std-Hyb`⟩ over time for a sample process. Normalization as described in Sup. Sec.S1.3 was performed for easier interpretation.

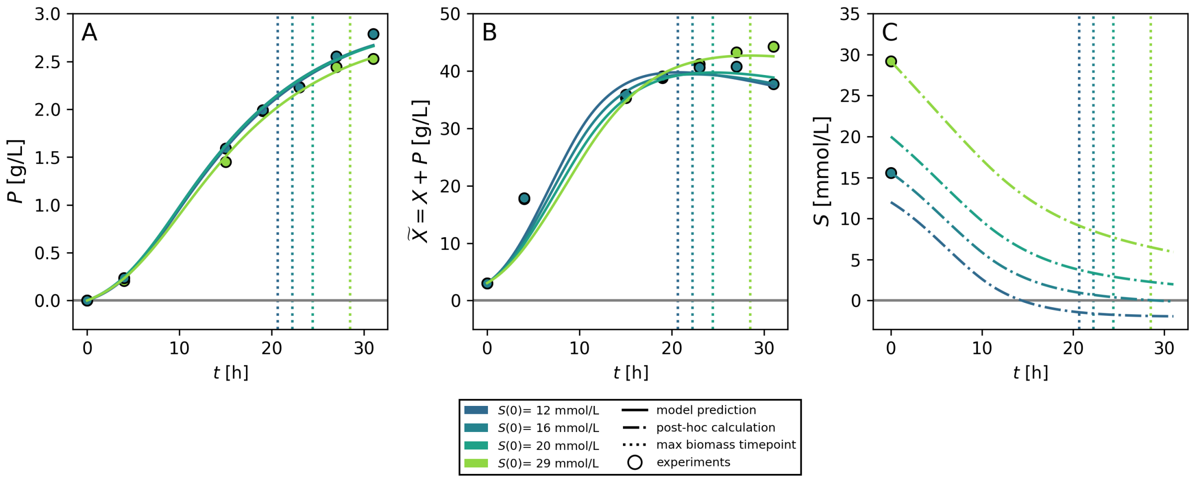


**Figure S6: SLIM optimization with `Std-Hyb`.** Panels **A** and **B** show predicted values for product ($P$) and apparent biomass ($\widetilde{X}$) for different initial sulfate concentrations. Panel **C** shows the estimated values of sulfate ($S$) after prediction from ⟨`Std-Hyb`⟩ via traditional FBA. It is visible that more sulfate is required for the predicted biomass than sulfate is available. The sulfate concentration cannot be implemented into ⟨`Std-Hyb`⟩ in a straightforward manner.

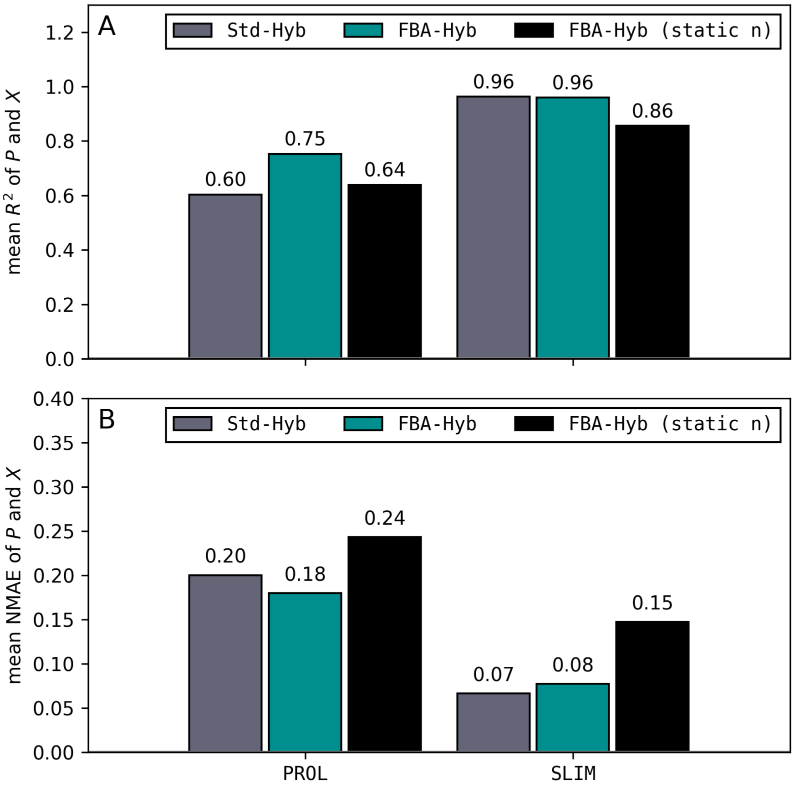


**Figure S7: LOPO CV results for static n** for both PROL (panel **A**) and SLIM (panel **B**) case studies. One can see that if we find a single, process condition-independent, value for $\mathbf{n}$ in the `FBA-Hyb` formulation, both $R^2$ and NMAE are worse than with a dynamic, process condition-dependent, prediction of $\mathbf{n}$. Interestingly, in the selected case studies, even the `Std-Hyb` outperforms the fixed-$\mathbf{n}$ `FBA-Hyb`. Methodologically, an `FBA-Hyb` with static $\mathbf{n}$ is very close to a dynamic FBA, the major difference being that in the former, $q_G$ is predicted by an ANN, whereas in the latter usually $q_G$ is estimated by a mechanistic, often Michaelis Menten-type equation.
